# Modulation of antigen discrimination by duration of immune contacts in a kinetic proofreading model of T cell activation with extreme statistics

**DOI:** 10.1101/2023.05.30.542789

**Authors:** Jonathan Morgan, Alan E. Lindsay

## Abstract

T cells form transient cell-to-cell contacts with antigen presenting cells (APCs) to facilitate surface interrogation by membrane bound T cell receptors (TCRs). Upon recognition of molecular signatures (antigen) of pathogen, T cells may initiate an adaptive immune response. The duration of the T cell/APC contact is observed to vary widely, yet it is unclear what constructive role, if any, such variations might play in immune signaling. Modeling efforts describing antigen discrimination often focus on steady-state approximations and do not account for the transient nature of cellular contacts. Within the framework of a kinetic proofreading (KP) mechanism, we develop a stochastic *First Receptor Activation Model* (FRAM) describing the likelihood that a productive immune signal is produced before the expiry of the contact. Through the use of extreme statistics, we characterize the probability that the first TCR triggering is induced by a rare agonist antigen and not by that of an abundant self-antigen. We show that defining positive immune outcomes as resilience to extreme statistics and sensitivity to rare events mitigates classic tradeoffs associated with KP. By choosing a sufficient number of KP steps, our model is able to yield single agonist sensitivity whilst remaining non-reactive to large populations of self antigen, even when self and agonist antigen are similar in dissociation rate to the TCR but differ largely in expression. Additionally, our model achieves high levels of accuracy even when agonist positive APCs encounters are rare. Finally, we discuss potential biological costs associated with high classification accuracy, particularly in challenging T cell environments.

**Author summary:** Physical contact between the T cell and antigen presenting cell (APC) is essential for productive immune signaling. Wide variations in this contact time have been observed yet little is known of mechanisms controlling this crucial timescale, nor how its duration may impact antigen discrimination. We develop and analyze a probabilistic mathematical model of T cell activation which combines kinetic proofreading (KP) with a finite contact duration. Our model is capable of suppressing large populations of self ligands while remaining sensitive to only a single agonist in T cell/APC cellular contacts. Additionally, we explored two challenging cases, one in which self and agonist antigen are similar and one in which agonist positive APCs are rare. We found that our model could overcome these environmental challenges by increasing the number of kinetic proofreading steps. Finally, we discuss the potential biological costs of achieving such accuracy. Our work demonstrates the extreme effectiveness of kinetic proofreading in a temporal context while also demonstrating the possible challenges in biological implementation of such a model.

## Introduction

### Background

T cells are immune cells that continuously search for molecular signatures (antigens) of pathogens and upon recognition, can initiate an adaptive immune response. When a T cell encounters an antigen presenting cell (APC), T cells recognize antigen through binding of their T cell receptors (TCR) to the peptide-MHC complex (pMHC) on the APC. For an efficient immune response, T cells must be able to recognize when an APC is agonist positive, where an APC displays foreign antigen indicating an immune response is appropriate, or agonist negative, where an APC displays only self antigen indicating no response is the appropriate action. This recognition is accomplished at the level of receptor/ligand interactions, where TCR/pMHC binding can result in the generation of an intracellular signal, such as Ca^2+^ influx, that may lead to T cell activation [1]. This process of antigen recognition is marked by rapid recognition speeds with single-molecule sensitivity to agonist ligands [2]. For example, CD4^+^ T cells can exhibit Ca^2+^ signaling when stimulated with a single molecule of foreign antigen [3, 4]. Similarly, CD8^+^ T cells have been shown to recognize as few as 3 molecules of a foreign antigen [5]. This single-digit molecule sensitivity is particularly remarkable given how short-lived TCR/pMHC interactions can be [6–8]. In addition, T cells can amplify small differences in antigen affinity into large differences in their responses [9–15]. The combination of these features of T cells have led many to characterize T cell antigen discrimination as being near-perfect [16–22], where the T cell is capable of recognizing agonist positive APCs (APCs with at least one agonist antigen on the surface) and simultaneously remain non-reactive to agonist negative APCs (APCs with no agonist presence) even when agonist populations are small, self antigen populations are large, and the distinguishing characteristic between agonist and self antigen is a slight difference in TCR affinity. Moreover, some studies have shown that TCRs may be more specifically tuned to the antigen dissociation rate, rather than the affinity [6, 11, 23–27]. This may indicate that T cells are specifically tuned to recognize agonist and self antigen primarily based on the antigen dissociation rate with the TCR.

### Kinetic proofreading and cellular contact times

The observed features of T cell antigen discrimination led McKeithan to introduce a kinetic proofreading (KP) model (Figure 1) for TCR activation. KP proposes that a productive immunological signal can arise after the completion of several intermediate conformational changes, all while being subject to TCR/pMHC disassociation that may eliminate progress towards activation [28]. KP has since become a prominent and highly studied conceptual framework for understanding antigen discrimination [29–34]. Steady state analysis of KP models shows that the mechanism is certainly capable of ligand discrimination, however, large increases in the number of intermediate states would increase the specificity of the model, but only at the expense of model sensitivity to agonists [4, 35], i.e., the KP mechanism could reproduce observed ligand discrimination only when the agonist population was sufficiently large. This is because the increase in intermediate bound states would yield an unacceptably low steady state fraction of activated TCRs when there were only few agonist antigen. This result calls into question the feasibility of KP models being the sole descriptor of the observed antigen discrimination in T cells [16–22, 36], since both, high specificity and sensitivity to small populations of agonists, have been observed in experiments. Nevertheless, despite the shortcomings of theoretical models, KP is supported by observations that show the antigen dissociation rate is a significant indicator of self or agonist antigen, and experimental work has continued to show results that indicate kinetic proofreading is involved in the antigen recognition process [34, 37, 38]. It should also be noted that the ligand discrimination observed in early experiments has also been called into question [39] and it has been hypothesized that T cell antigen discrimination is not as “good” as previously believed, potentially restoring the plausibility of KP as a model for antigen discrimination.

**Fig 1.**
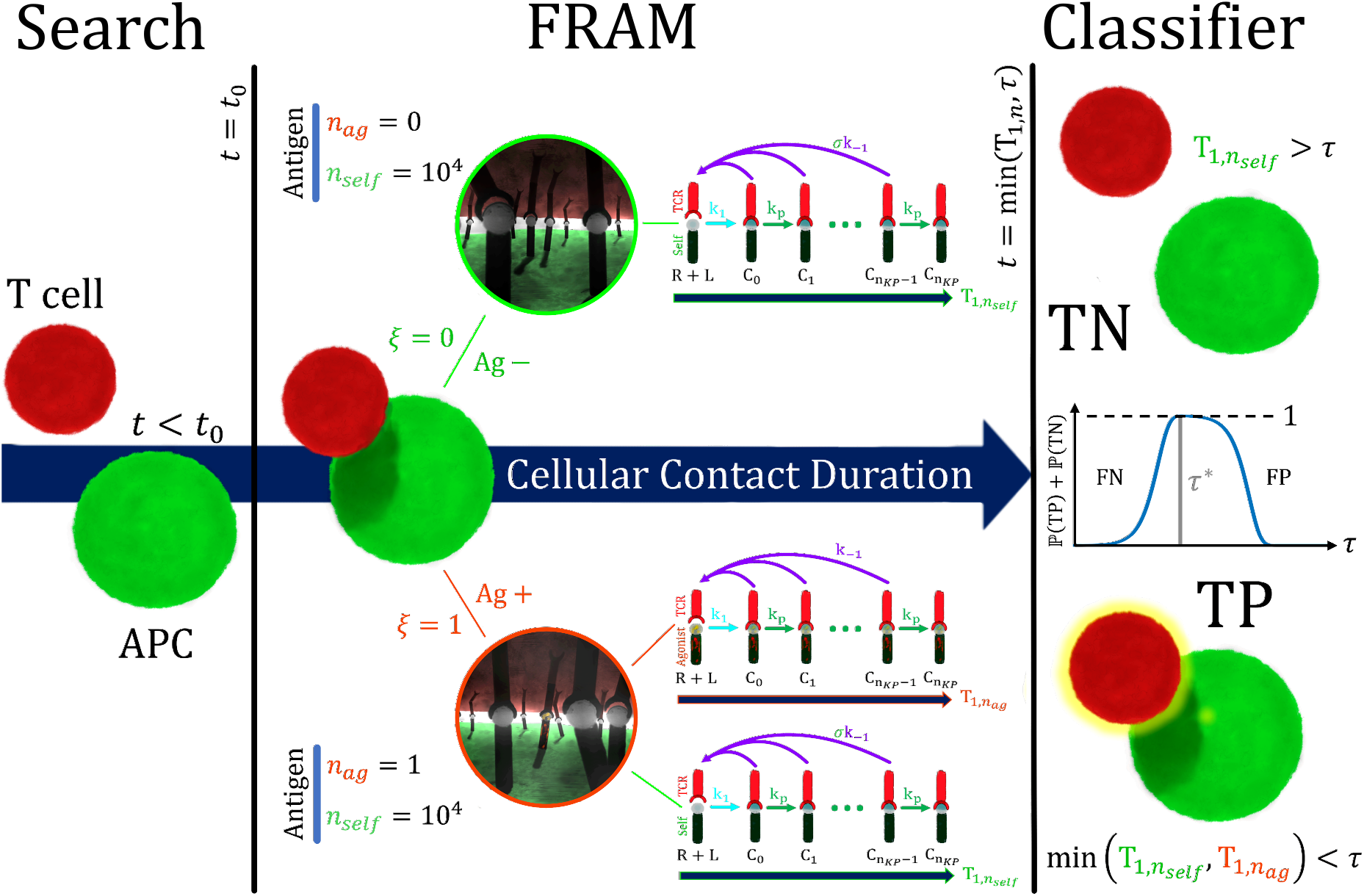
The First Receptor Activation Model (FRAM) as a classifier of APC status *ξ∈ {*0, 1*}*. The T cell and APC form a cellular contact at *t* = *t*_0_. An APC is agonist positive (*ξ*= 1) with probability ℙ (*ξ*= 1) = *ρ*_*ag*_ and agonist negative (*ξ*= 0) with probability ℙ (*ξ*= 0) = 1 *− ρ*_*ag*_. If *ξ*= 1 then there is *n*_*ag*_ = 1 agonist antigen and *n*_*self*_ = 10^4^ self antigen. Activation is the event 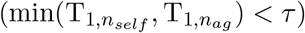 which results in a true positive (TP) classification. If *ξ*= 0 then correct classification occurs when none of the self antigen activate 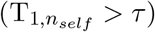 which results in a true negative (TN). We measure the accuracy (ℙ (TP) + ℙ (TN)) of the FRAM as an APC classifier with respect to the cellular contact duration (*τ*). Cellular contacts of too short duration result in a high false negative rate while overly long contacts return a high false positive probability, with both scenarios reducing accuracy. Decision accuracy is maximized at fixed contact duration (*τ* ^*∗*^).

It has been estimated that APCs encounter 500-5000 T cells every hour (dendritic cells) [40–42], suggesting many T cell/APC contacts are formed over short periods in time. Experimental work has shown evidence that the duration of these contacts may be significant in antigen recognition [43–49] and some studies have classified the engagement period into distinct phases based on cellular contact times [50–55]. The first (phase I) of these periods may involve multiple, short-duration, transient encounters between T cells and APCs which persist until a certain threshold of accumulated signal followed by a second period (phase II) in which a long, stable contact forms. In the context of a KP mechanism of activation, it is clear that a sufficiently long-lasting cellular contact is required in order for a productive immune signal to be generated and so early termination of the cellular contact may hypothetically prevent T cell activation [56]. In addition, early termination of a T cell/APC contact could prevent the transition from a short transient contact to a more stable contact. Together, this suggests that the duration of the T cell/APC contact may have an important role in antigen discrimination.

When the duration of the T cell/APC contact is explicitly modeled, the previously employed steady-state analysis may not be appropriate [28, 30, 36, 57, 58]. For example, if the cellular contact duration is shorter than the timescale of relaxation to the steady state, the equilibrium configuration may not be attained. Similarly, in scenarios where activation is defined by an accumulation of productive signal, integration over periods of non-equilibrium behavior results in a non-trivial dependence of contact duration [59]. Finally, in rebinding models where multiple short TCR/antigen engagements [7, 17, 19, 35] can lead to rare TCR triggering events, the role of a finite cellular contact duration may be a significant factor in modeling immunological outcomes.

### Summary of results

In this paper, we develop a *First Receptor Activation Model* (FRAM) which describes T cell activation as the event when any one receptor is triggered by an antigen ligand before the cellular contact expires. For a population of APCs where a fraction (*ρ*_*ag*_) are agonist positive, we determine the accuracy of a T cell as an APC classifier, which is the probability Γ that a T cell will correctly classify a randomly chosen member of this population. Mathematically, this involves calculating certain extreme statistics [60–63] due to the fact that while a single T cell/APC interaction generates numerous independent TCR/pMHC reaction pairs, it is the fastest of these that will set the activation time. A positive outcome requires our model to be reactive to a small number of agonist antigen while non-reactive during numerous interactions with abundant self-antigen.

We observe that the accuracy of our model varies considerably with contact duration *τ* and demonstrate that there often exists a window of time such that Γ *≈* 1, i.e. the accuracy is near perfect. We find that regions of high accuracy expand/contract as the number *n*_*KP*_ of KP states increases/decreases. A similar relationship is observed with respect to variations in the ratio *σ* between dissociation rates of self and agonist antigen (Figure 1. Specifically, classification accuracy is higher at larger *σ* values and decreases as *σ →* 1^+^. We show that our model can achieve near-perfect classification accuracy for just a single agonist antigen *n*_*ag*_ = 1, *n*_*self*_ = 10^4^ self antigen, and a relative dissociation factor *σ* = 2. This demonstrates that by increasing the number of kinetic proofreading steps the FRAM is able to overcome the challenge of remaining sensitive to a single angonist antigen, and suppressing receptor triggering for a large self population of antigen, even when self and agonist antigen appear similar in the lens of a KP mechanism.

The proportion *ρ*_*ag*_ of agonist positive APCs influences the difficulty of the classification task. This is especially so when *ρ*_*ag*_ is small since this means most cellular contacts are made of agonist negative cells, and the suppression of large self antigen populations becomes more significant. When agonist positive cells are rare, we found that shorter-duration contacts decrease the likelihood that a T cell activation is a result of a false positive. This demonstrates that the agonist positive prevalence may influence optimal cellular contact durations. However, our results also demonstrate that a sufficient number of KP steps could overcome this challenge by maximizing the accuracy of the T cell for all *ρ*_*ag*_ and effectively making the optimal cellular contact durations independent of the agonist positive prevalence.

Lastly, we quantify a potential biological constraint in the FRAM. We assume that forward KP reactions involve events with an energy cost [1, 28, 64–66], which we call *futile* reactions if the TCR/antigen dissociate before TCR activation. As mentioned, our results show that increasing *n*_*KP*_ could overcome the challenges of similar self/agonist ligands, large differences in self/agonist expression, and rare agonist positive APCs. However, we show that the increased number of KP steps results in either longer-lasting cellular contacts or faster KP rates, both of which cause an increased number of futile reactions. We conclude that accurate antigen discrimination may come with associated energetic costs.

### Model

We consider a population of APCs, each of which is agonist positive (*ξ*= 1) with probability ℙ (*ξ*= 1) = *ρ*_*ag*_ or agonist negative (*ξ*= 0) with probability ℙ (*ξ*= 0) = 1 *− ρ*_*ag*_. In the case *ξ*= 1 there are *n*_*self*_ = 10^4^ self antigen and *n*_*ag*_ = 1 agonist antigen in any T cell/APC contact while for *ξ*= 0 there are *n*_*self*_ = 10^4^ self antigen and *n*_*ag*_ = 0 agonist antigen in any contact. We assume that during T cell interrogation of an APC surface, the TCR density is sufficient to engage all antigen in the contact. Each TCR/antigen pair undergoes dynamics according to the KP mechanism Figure 1 and the first receptor activation model (FRAM) defines T cell activation as the event when any TCR reaches the signaling state 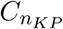.

We define 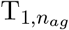 to be the signaling time of an agonist antigen and agonist activation to be the event 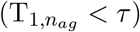. Self activation of the T cell occurs when one of the *n*_*self*_ = 10^4^ self antigen [14] activate a TCR before the cellular contact expires 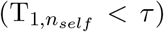. We emphasize that the self activation time is 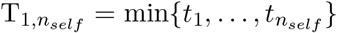 where *t*_*i*_ is the activation time of the *i*^*th*^ signaling pair and is hence an example of an extreme statistic [60–63, 67]. Extreme events are typically very fast, yet positive immunological outcomes will require that 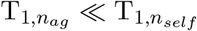.

We derive an analytical expression for the first passage time 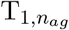 of TCR activation in the case of a single TCR/pMHC pair (supplement). For a large population of ligands and receptors, we approximate the distribution for the extreme statistic 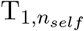 by introducing a system of ordinary differential equations (supplement). In addition, we connect the limiting case *n*_*self*_ *→ ∞* with recent results of extreme statistic for continuous time Markov chains with discrete states [67]. Specifically, as *n*_*self*_ *→ ∞*, we confirm (supplement) the limiting behavior

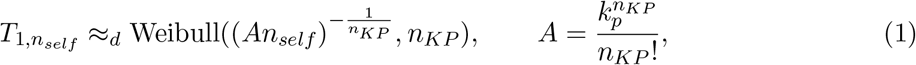

where a random variable *X ≥* 0 has a *Weibull* distribution with scale parameter *λ >* 0 and shape parameter 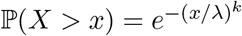. In such a case, we define *X* = Weibull(*λ, k*).

In the event that *ξ*= 1, both activation cases result in a true positive since the T cell activated and was in contact with an agonist positive APC. The T cell does not activate when none of the *n*_*self*_ = 10^4^ self antigen activate a TCR before the cellular contact expires 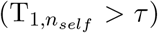. When *ξ*= 0, this results in a true negative.

We utilize the FRAM as an APC classifier, where the task is to correctly identify agonist positive and agonist negative APCs. In our model, a T cell makes contact with an APC at time *t* = 0 and remains in contact till the expiry time *t* = *τ*. A true positive classification occurs when an agonist positive APC is activated before contact expiry (*ξ*= 1 and min 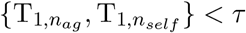). A true negative classification occurs when a T cell is in contact with an agonist negative APC and the T cell does not activate before the expiration of the cellular contact (*ξ*= 0 and 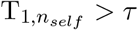). Mathematically, the four outcomes of the FRAM have probabilities

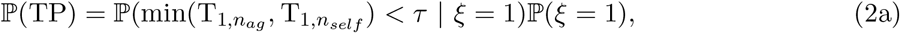

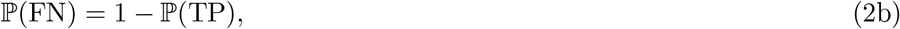

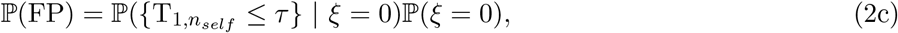

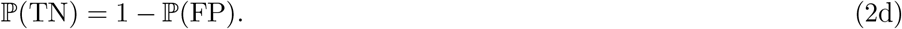

The main parameters of interest are *{τ, ρ*_*ag*_, *n*_*KP*_, *σ}*, where *τ >* 0 is the expiry time, *ρ*_*ag*_ *∈* (0, 1) is the agonist positive prevalence, *n*_*KP*_ is the number of KP steps, and *σ >* 1 is the ratio of TCR/antigen dissociation rates between self and agonist antigen which is referenced in previous experimental and theoretical works [28, 30, 59, 65, 68]. Finally, we make the assumption that the first receptor activation by self antigen is independent of the first receptor activation by agonist antigen, i.e.,

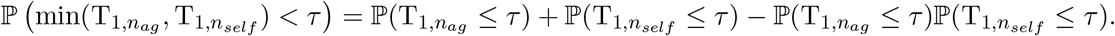

Since we are modeling the self and agonist situations separately in the agonist positive T cell/APC contacts, then independence is guaranteed. However, this is clearly an approximation since we do not account for possible interactions, or interference, between self and agonist ligands.

We explore the properties of the FRAM by quantifying the T cell accuracy

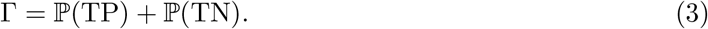

The accuracy shares similarities with some earlier measures of sensitivity and specificity in kinetic proofreading models [30], however, there are two main differences. First, the measure Γ combines both sensitivity and specificity (each can be individually described through P(TP) and P(TN) into a single measurement. The second is that we are applying the measure at the population level of T cells, or more accurately, a population of T cell/APC contacts, and not at the level of individual receptors/ligand interactions Figure 1.

A list of parameters and variables together with meanings and ranges are given in Table 1. The derivation and numerical validation of the probabilities (2) together with the formula for the accuracy (3) are given in the supplement. The focus of the remainder of the article is analyzing variations in the accuracy Γ with the parameters *{τ, ρ*_*ag*_, *n*_*KP*_, *σ}*.

**Table 1.**
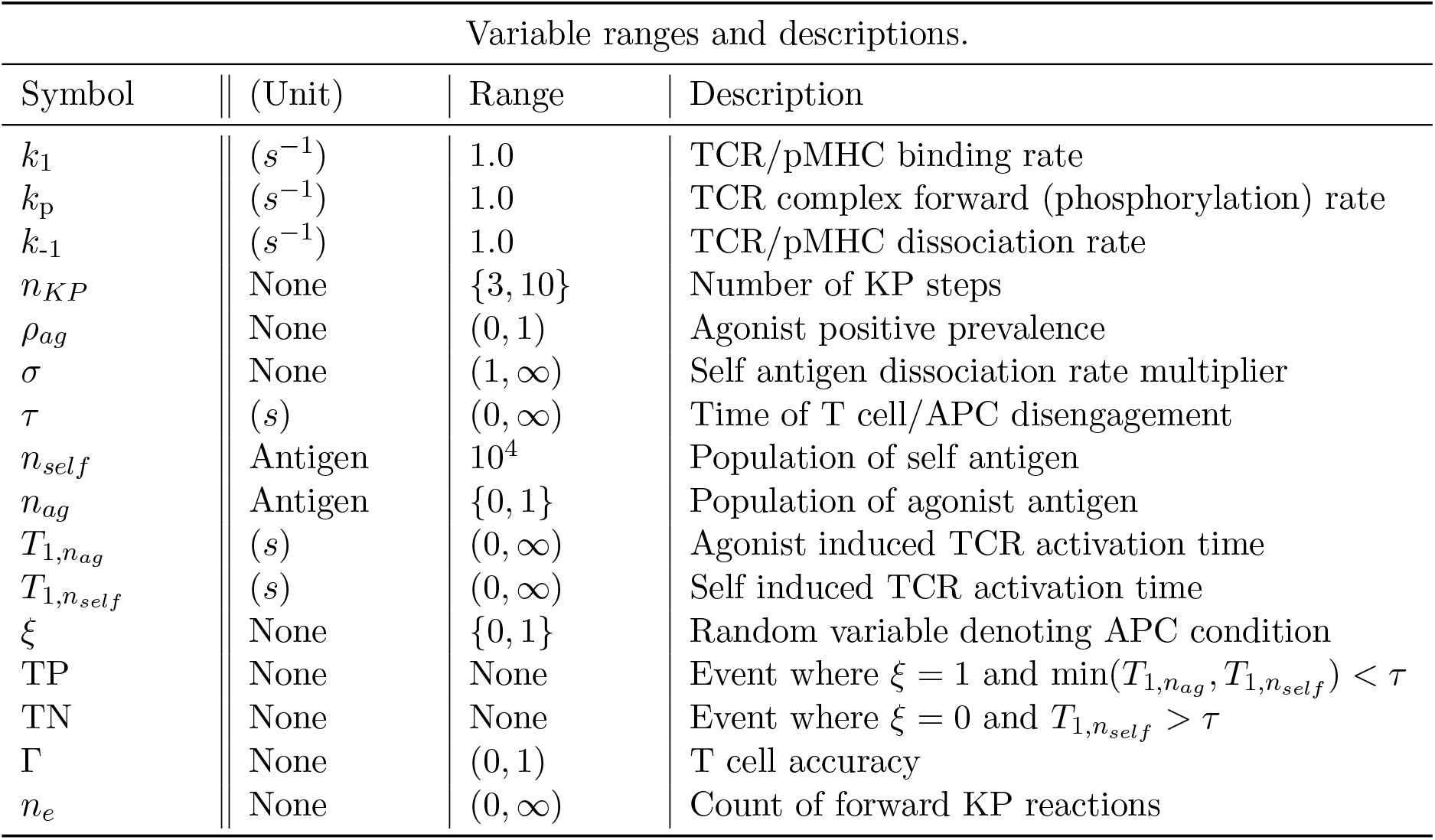
A list of model parameters and variables with their default values or ranges (unless otherwise specified).

| Variable ranges and descriptions. |  |  |  |
| --- | --- | --- | --- |
| Symbol | (Unit) | Range | Description |
| $k_1$ | $(s^{-1})$ | 1.0 | TCR/pMHC binding rate |
| $k_p$ | $(s^{-1})$ | 1.0 | TCR complex forward (phosphorylation) rate |
| $k_{-1}$ | $(s^{-1})$ | 1.0 | TCR/pMHC dissociation rate |
| $n_{KP}$ | None | $\{3, 10\}$ | Number of KP steps |
| $\rho_{ag}$ | None | $(0, 1)$ | Agonist positive prevalence |
| $\sigma$ | None | $(1, \infty)$ | Self antigen dissociation rate multiplier |
| $\tau$ | $(s)$ | $(0, \infty)$ | Time of T cell/APC disengagement |
| $n_{self}$ | Antigen | $10^4$ | Population of self antigen |
| $n_{ag}$ | Antigen | $\{0, 1\}$ | Population of agonist antigen |
| $T_{1,n_{ag}}$ | $(s)$ | $(0, \infty)$ | Agonist induced TCR activation time |
| $T_{1,n_{self}}$ | $(s)$ | $(0, \infty)$ | Self induced TCR activation time |
| $\xi$ | None | $\{0, 1\}$ | Random variable denoting APC condition |
| TP | None | None | Event where $\xi = 1$ and $\min(T_{1,n_{ag}}, T_{1,n_{self}}) < \tau$ |
| TN | None | None | Event where $\xi = 0$ and $T_{1,n_{self}} > \tau$ |
| $\Gamma$ | None | $(0, 1)$ | T cell accuracy |
| $n_e$ | None | $(0, \infty)$ | Count of forward KP reactions |

## Results

### Influence of contact duration on T cell accuracy

We explored the influence of the T cell/APC contact duration on T cell accuracy by computing Γ(*τ*) over a range of *τ* for several values of *σ* and *n*_*KP*_ (Figure 2A-C). When *ρ*_*ag*_ = 0.5, we have the lower bound Γ ⪆ 0.5 indicating poor APC classification accuracy (i.e. ℙ (TP) *≈* ℙ (FP)). If *n*_*KP*_ and *σ* are insufficiently large, the FRAM cannot accurately classify APC contacts for any *τ* (*σ* = 10^2^ in Figure 2A). However, in the cases where *σ* is sufficiently large (*σ ≥* 10^4^ in Figure 2A) and/or the number of proofreading steps is sufficiently large (*σ ≥* 10 in Figure 2B), a range of contact times exists where Γ(*τ*) *≈* 1 (*τ ∈* [10^4^, 10^5^] and *σ* = 15 in Figure 2B).

**Fig 2.**
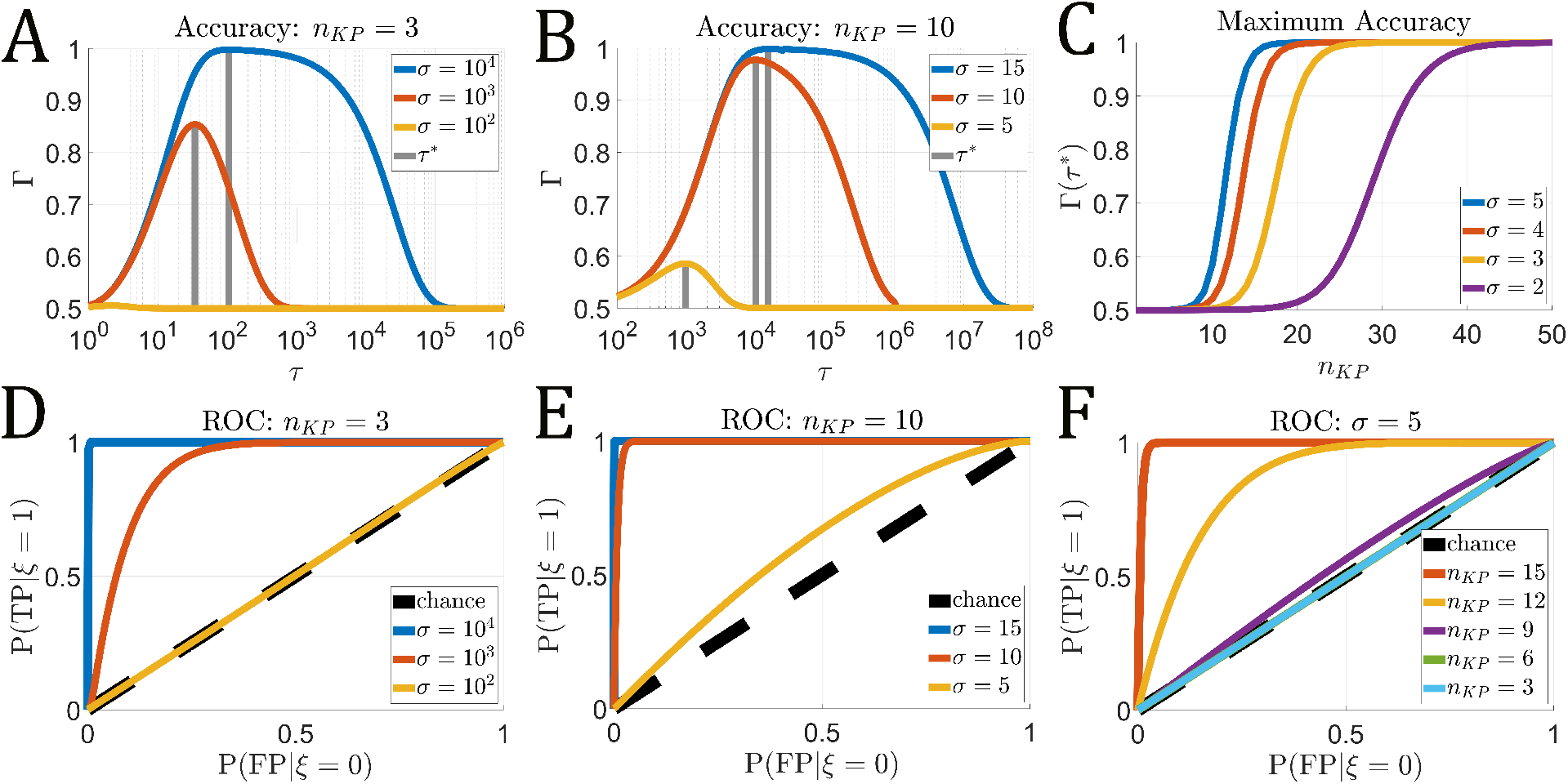
T cell accuracy as contact duration *τ* varies. Kinetic proofreading parameters given in Table 1. **A–B** The T cell classification accuracy Γ(*τ*) for *n*_*KP*_ = 3 (**A**) and *n*_*KP*_ = 10 (**B**). For each value of *σ*, the time *τ* ^*∗*^ of maximum accuracy is highlighted (gray vertical lines). **C** The maximum accuracy Γ(*τ* ^*∗*^) as *n*_*KP*_ varies for values *σ* = *{*2, 3, 4, 5*}*. **D–F** Receiver operating characteristic (ROC) for *n*_*KP*_ = 3 (**D**) and *n*_*KP*_ = 10 (**E**) at various *σ* values. **F** The ROC for *σ* = 5 and varied *n*_*KP*_.

To identify scenarios where effective APC classification is achieved, we determine the maximum accuracy over a range of contact durations,

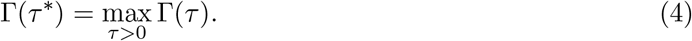

By increasing *n*_*KP*_, we find that one can attain a near-perfect classification accuracy at the optimal T cell/APC contact duration (Γ(*τ* ^*∗*^) *≈* 1), even in cases where agonist and self antigen only differ slightly in dissociation rate but differ largely in expression. Since Γ(*τ* ^*∗*^) = 1 if and only if ℙ (TP|*ξ*= 1; *τ* ^*∗*^) = 1 and ℙ (TN|*ξ*= 0; *τ* ^*∗*^) = 1, this indicates that the T cell always activates when in contact with an agonist positive APC and never activates when in contact with an agonist negative APC (Figure 2C).

The receiver operating characteristic (ROC) is plotted in Figure 2D-F and allows for a more nuanced consideration of classification outcomes over the parameter *τ*. An indicator of a classifier’s performance is the area under the curve (AUC). We see that the maximum AUC (AUC *≈* 1) can be achieved by increasing *σ* (Figure 2D and E) and/or increasing *n*_*KP*_ (Figure 2F), which indicates the model is capable of identifying agonist positive and agonist negative contacts with near-perfect accuracy. The worst-case AUC for the FRAM is when ℙ(TP|*ξ*= 1) *≈* ℙ(FP|*ξ*= 0) for all *τ*. This equality also yields ℙ(*ξ*= 0|T_1,*n*_ *< τ*) = *ρ*_*ag*_ and ℙ(*ξ*= 1|T_1,*n*_ *< τ*) = 1 *− ρ*_*ag*_ for all *τ*. This means that in these cases, the FRAM as a classifier can do no better than classifying whichever population is in the majority. We call this observation the baseline and discuss this more in-depth in the next section. Similar to our result in Figure 2C, we show that increasing the number of kinetic proofreading steps allows for arbitrary increases in the AUC, even when *σ* is small (Figure 2F).

### The effects of an agonist positive prevalence

Previously we assumed an equal proportion of agonist positive and negative APCs (*ρ*_*ag*_ = 0.5). Variations in this prevalence can lead to significant changes in the accuracy and optimal cellular contact duration, as we now demonstrate by varying *ρ*_*ag*_ between 0 and 1.

In Figure 3 A–C we plot the optimal accuracy Γ(*τ* ^*∗*^) against *ρ*_*ag*_ for several values of *n*_*KP*_ and *σ*. In Figure 3A, we observe at the value of *σ* = 10^4^ that Γ *≈* 1 for all *ρ*_*ag*_, indicating accurate classification. As *σ* decreases and/or *n*_*KP*_ is too small, we find that the FRAM is incapable of accurately identifying the agonist positive or agonist negative cells for small or large *ρ*_*ag*_, respectively. In these cases, the *baseline* limiting case for the accuracy becomes

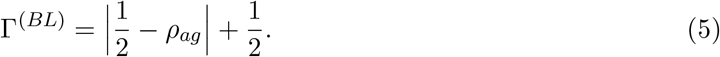

Equation (5) reflects the probability of an accurate T cell response with the strategy of always activating when 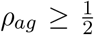 and never activating when 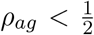. In Figure 3B we show results for *n*_*KP*_ = 10 and observe higher accuracy at smaller values of *σ* and over larger ranges of *ρ*_*ag*_ when compared to *n*_*KP*_ = 3. In Figure 3C we observe that even for small *σ*, perfect accuracy can be obtained for all *ρ*_*ag*_ by increasing *n*_*KP*_.

**Fig 3.**
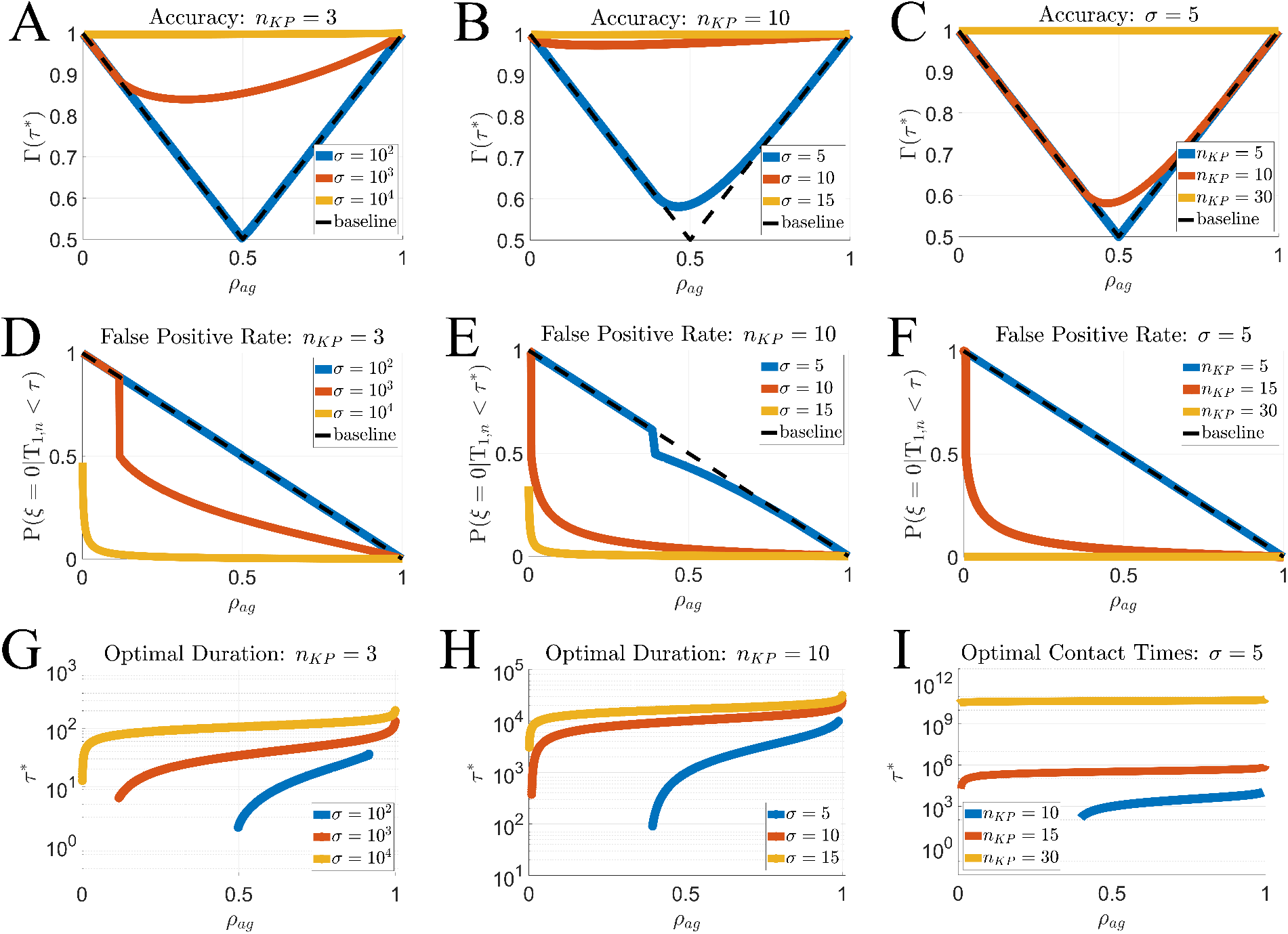
Effect of agonist positive prevalence (*ρ*_*ag*_) on T cell accuracy (Γ(*τ* ^*∗*^)) and false positive rate at the optimal contact duration (6). **A–C** Accuracy Γ(*τ* ^*∗*^) versus *ρ*_*ag*_ for *n*_*KP*_ = 3 (**A**), *n*_*KP*_ = 10 **B**, and varied *n*_*KP*_ with *σ* = 5 (**C**). The baseline (black dashed line) shows the worst accuracy a model can accomplish given that the optimal cellular contact duration is chosen. **D–F** False positive rate (6) at the maximizing cellular contact time (*τ* ^*∗*^). **G–I** Effect of agonist positive prevalence on the optimal cellular contact duration. Curve endpoints indicate where the FRAM reduces to baseline, i.e., *τ* ^*∗*^ *→* 0 if *ρ*_*ag*_ *<* 0.5 or *τ* ^*∗*^ *→ ∞* if *ρ*_*ag*_ *>* 0.5.

When agonist positive contacts are rare (0 *< ρ*_*ag*_ *≪* 1), accurate classification becomes more challenging and the risk of false positives is greater. We use the outcome probabilities and determine the false positive rate ℙ (*ξ*= 0|T_1,*n*_ *< τ* ^*∗*^)

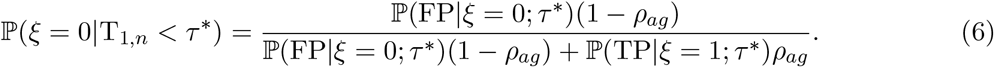

which is defined as In equation (6), evaluation is at the optimal cellular contact duration *τ* ^*∗*^ and T_1,*n*_ is the activation time of the FRAM with an unknown APC condition. As *ρ*_*ag*_ decreases, the probability that any T cell activation is a false positive increases (Figure 3 D–F). Again, as accurate APC identification becomes too difficult, we identify convergence to the baseline case

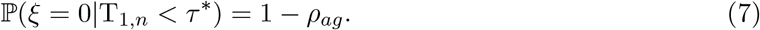

In Figure 3 D and E, we demonstrate that increasing *σ* can greatly reduce the likelihood of false positives. However, even the cases that were sufficient to yield nearly 100% accuracy (*σ* = 15 in Figure 2B), still yield a high false positive rate when *ρ*_*ag*_ is small (*σ* = 15 in Figure 3E). However, in Figure 3F) we choose *σ* = 5 and show that increasing *n*_*KP*_ can completely inhibit the false positive rate (P(*ξ*= 0|T_1,*n*_ *< τ* ^*∗*^) *≈* 0) for nearly all *ρ*_*ag*_.

In the FRAM, a decrease in cellular contact duration serves to decrease the probability of activation in any T cell/APC encounter. The curves in Figure 3G–I represent ranges of *ρ*_*ag*_ where the FRAM has some ability to classify both APC conditions, not just the majority. The endpoints of these curves represent the transition to the baseline accuracy (4) where *τ* ^*∗*^ *→ ∞* if 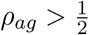, and *τ* ^*∗*^ *→* 0 if 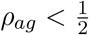. As *ρ*_*ag*_ decreases, the false positive rate increases. In light of this, we observe that *τ* ^*∗*^ decreases with *ρ*_*ag*_ (Figure 3G–I). Increasing *σ* (Figure 3G and H) reduces the influence of *ρ*_*ag*_ on the cellular contact duration and decreases the range of *ρ*_*ag*_ where accuracy reduces to the baseline (4). In Figure 3I, we note that the contact duration becomes nearly independent of *ρ*_*ag*_ as *n*_*KP*_ is increased.

Taken together, these results suggest that the accuracy is independent of the agonist positive prevalence when *σ* and/or *n*_*KP*_ is sufficiently high. Otherwise, small/large *ρ*_*ag*_ can decrease/increase the optimal cellular contact durations and decrease the ability of the FRAM to recognize rare APC conditions without a significant risk of false positives/negatives. We also note an asymmetry in each panel of Figure 3A–C and Figure 3G–I where the accuracy Γ(*τ* ^*∗*^) more quickly reduces to the baseline (4) at smaller values of *ρ*_*ag*_. This demonstrates that the false positive rate may be more problematic than the false negative rate for T cell classification accuracy.

### High decision accuracy comes at the cost of increased number of energy utilizing reactions

Theoretical works have hypothesized that kinetic proofreading involves energy-consuming reactions [29–33], for example phosphorylations of the TCR complex following TCR/pMHC binding [1, 38, 69]. To estimate such a cost, we approximate the mean number of futile reactions *n*_*e*_, i.e., the forward reactions in the KP mechanism that do not result in TCR activation over a contact duration *τ*. We estimate this quantity by deriving and solving an associated mass action ODE system (supplement). We do not count the initial binding event so that the cost to reach the state *C*_*i*_ from state *C*_0_ is *i* forward reactions. A single TCR/pMHC complex that reaches the *i*^*th*^ bound state in the KP mechanism (Figure 1) before dissociating results in *i* futile reactions (*i < n*_*KP*_).

In Figure 4 we observe the relationship between T cell accuracy and energetic reactions by plotting a scaled accuracy 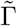 defined as

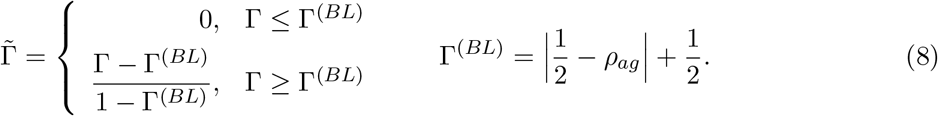

High T cell accuracy(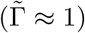) is associated with a large number of futile reactions, particularly when *σ* and *ρ*_*ag*_ are small. As *σ* and/or *ρ*_*ag*_ decreases (Figure 4 columns), a larger *n*_*KP*_ is needed to achieve moderate or large temporal regions of high accuracy. The increased energy requirement at smaller *σ* and *ρ*_*ag*_ values is a result of the increase in the number of KP steps. As *n*_*KP*_ increases, a longer contact duration is necessary to capture the first passage activation of an agonist antigen, i.e., observing a true positive activation. This time constraint can be reduced by increasing the KP rates of the FRAM (supplement). However, if the KP rates are scaled equally, then the number of futile reactions does not change (supplement).

**Fig 4.**
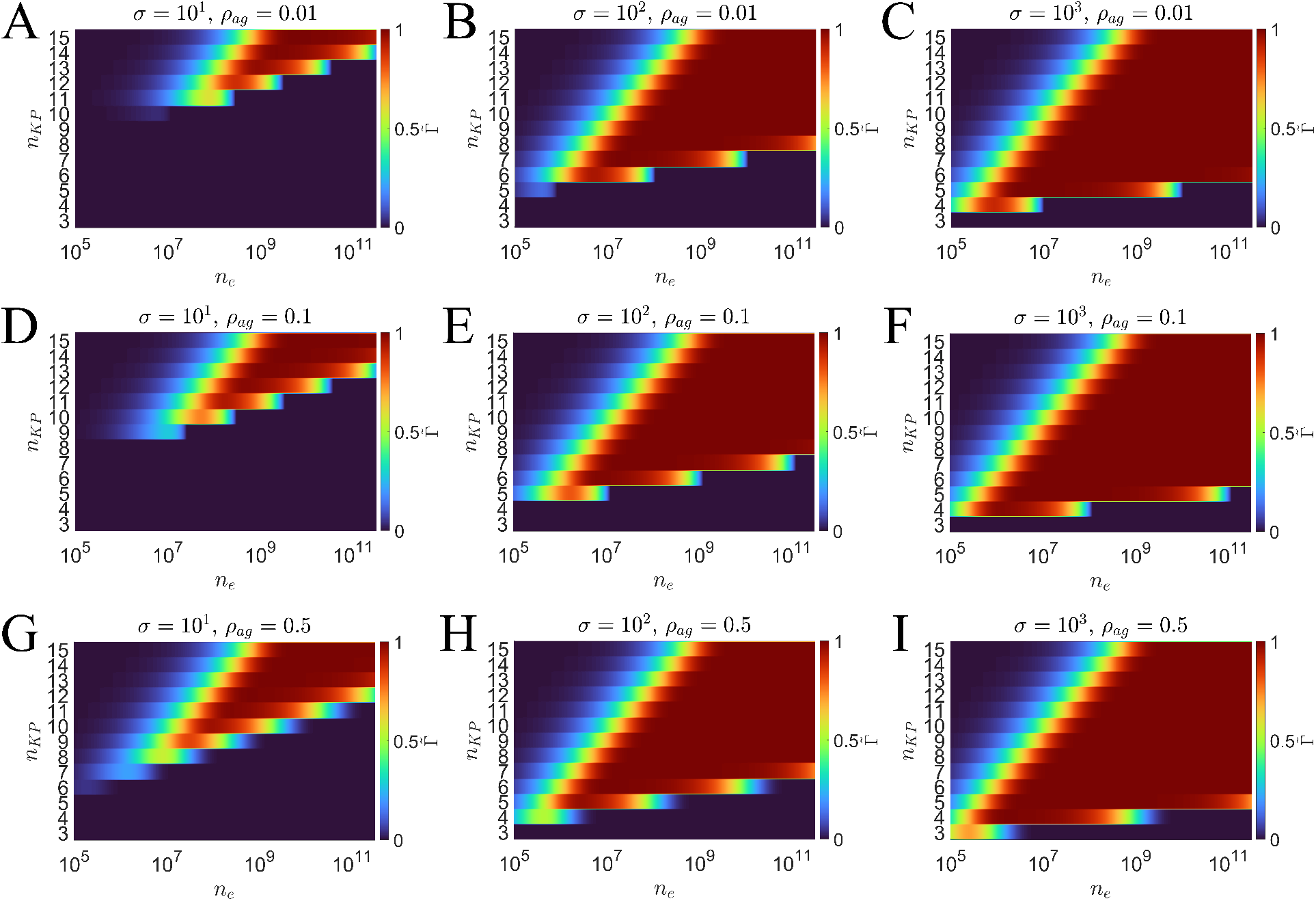
Plots showing the classification accuracy in terms of the number of energy utilizing reactions (*n*_*e*_) and *n*_*KP*_. The scaled accuracy 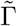 is such that 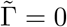 is Γ *≤* Γ^(*BL*)^ and 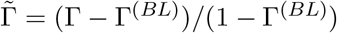. The left, center, and right column of plots show results for *σ* = 10, *σ* = 10^2^, *σ* = 10^3^, respectively. Each top, center, and bottom row shows results for *ρ*_*ag*_ = 0.01, *ρ*_*ag*_ = 0.1, and *ρ*_*ag*_ = 0.5

## Discussion

### Summary of results

We developed a first receptor activation model (FRAM) to evaluate the T cell as an APC classifier with a finite decision time. We used mathematical analysis of extreme statistics to determine the probabilities of T cell activation by either a single agonist antigen or numerous self-antigen. We evaluated the model in a challenging environment such as when self and agonist antigen are similar in KP properties but differ largely in expression or when agonist positive APCs are rare in a population. We used the accuracy (3) to measure T cell activation outcomes over a range of cellular contact durations and environmental conditions in order to investigate the FRAM as a classifier of APC agonist status.

We found that a high classification accuracy can be achieved over a large window of cellular contact times (Figure 2) given a sufficiently large *n*_*KP*_ (kinetic proofreading steps) and/or *σ* (ratio of antigen disassociation rates). Outside this window, poor accuracy can arise from contacts that are either too short or too long due to false negatives and positives, respectively. In addition, our results showed that the FRAM could overcome the challenge of similar agonist/self antigen ligands (small *σ*) and large disparities in self/agonist expression (*n*_*ag*_ = 1 and *n*_*self*_ = 10^4^) by sufficiently increasing the number of KP steps (Figure 2C,F).

Accurate classification is more challenging when *ρ*_*ag*_ is small/large due to a higher false positive/negative rate. Additionally, we found that the agonist positive prevalence can influence the optimal contact duration. When the agonist positive prevalence is small, the false positive rate increases (Figure 3D–F) which yields shorter optimal contact durations(Figure 3G–I), since decreasing the contact duration reduces the probability of T cell activation. When the agonist positive prevalence is large, the false negative rate increases which yields longer optimal contact durations, since increasing the cellular contact time increases the probability of activation. Additionally, we found that the accuracy of the FRAM is effectively independent of the agonist positive prevalence when *n*_*KP*_ is sufficiently large (Figure 3C,F,I), even when *σ* is small. This demonstrates that the FRAM can simultaneously overcome the challenge of similar self/agonist ligand as well small agonist positive prevalence by sufficiently increasing the number of KP steps.

Lastly, we quantified the cost in achieving high classification accuracy by considering the number of futile reactions in the FRAM. We found that for smaller *ρ*_*ag*_ and/or *σ* values, more futile reactions are necessary to effectively recognize both agonist positive and agonist negative cells (Figure 4). The primary contributor to the number of futile reactions is the suppression of a large self antigen population in the kinetic proofreading mechanism. When *σ* is large (Figure 4C,F,I), this suppression is achieved with a smaller number of kinetic proofreading steps, thus reducing the number of futile reactions. When *ρ*_*ag*_ is large (Figure 4G–I), the FRAM is capable of achieving good classification accuracy with smaller numbers of *n*_*KP*_ since the false positive risk is low. We found that the most difficult case is when both, *ρ*_*ag*_ and *σ* are small Figure 4A. In this case, a large *n*_*KP*_ is necessary to successfully identify the rare agonist positive cells while remaining insensitive to the large proportion of agonist negative cells. This has the added effect of requiring a large number of futile reactions, since most encounters are with an agonist negative APC and the FRAM must be able to suppress receptor triggering from the numerous self antigen in each of these encounters.

### Conclusions

We found that the FRAM is capable of high classification accuracy when agonist and self antigen are similar and differ largely in expression within a T cell/APC contact (Figure 2C and F). In addition, we found that our model could achieve this high accuracy even when agonist positive APCs are rare (Figure 3 C,F,I). This suggests that in terms of first passage times of agonist and self antigen, kinetic proofreading is more capable of antigen discrimination than what has been observed in T cell experiments [9]. Furthermore, unlike previous works [28, 30, 59], the FRAM yields no trade-off between remaining sensitive to agonists and being able to distinguish between agonist and self antigen populations.

Our results indicate that while the FRAM is capable of high levels of accuracy, it may be associated with certain costs, or biological constraints. We showed that when agonist and self antigen are more alike (smaller *σ*), a larger *n*_*KP*_ is needed (Figure 2C), which increases the cellular contact duration necessary to capture the first passage activation of an agonist antigen (supplement). Additionally, we found that when agonist positive cells are rare, more *n*_*KP*_ are necessary to successfully suppress the false positive rate (Figure 3D–F), which again has the effect of increasing the optimal cellular contact duration (Figure 3I). This may suggest that the cellular contact duration could act as a cost for accurate T cell responses.

From a mathematical perspective, we can scale the cellular contact durations to any value by increasing the rates of the model (supplement). However, this can yield large reaction rates (*k*_*p*_ *>* 10^9^*s*^*−*1^ in Figure 3I) which may also have a biological constraint. We found that this method of scaling has no influence on the number of futile reactions (supplement), which are reactions that may require energy consumption. Just as with the cellular contact duration, we found that more futile reactions are necessary for accurate APC recognition as *n*_*KP*_ increases (Figure 4). Hence, our results also demonstrate how energetic costs may act as a constraint in T cell antigen discrimination. These potential costs may provide insight for observations in biological experiments in which large discrepancies are observed in dissociation rates between agonist and self antigen [68]. Additionally, this may further support the idea that kinetic proofreading alone is not sufficient to explain some of the observations in previous T cell antigen discrimination experiments [9–15, 36].

In conclusion, we have shown how viewing the classic kinetic proofreading mechanism through resilience to extreme statistics (self activation) offers a different perspective on the problem of antigen discrimination. By modeling T cell activation as a first passage time describing receptor triggering by agonist or self antigen in the kinetic proofreading mechanism, we were able to show that the FRAM was capable of near perfect accuracy in challenging environmental conditions. We also showed several potential costs that may act as biological constraints to high accuracy in the T cell environment. Our hope is that the simplicity of this model yields a foundation from which more complexity can be built with respect to observed T cell characteristics, such as Ca^2+^ signaling and microvilli structures.

## Notes

### Competing Interest Statement

The authors have declared no competing interest.

### Summary of Updates

Previous upload was old version

## References

1. Courtney AH, Lo WL, Weiss A. TCR signaling: mechanisms of initiation and propagation. Trends in biochemical sciences. 2018;43(2):108–123.

2. Huse M, Klein LO, Girvin AT, Faraj JM, Li QJ, Kuhns MS, et al. Spatial and Temporal Dynamics of T Cell Receptor Signaling with a Photoactivatable Agonist. Immunity. 2007;27(1):76–88.

3. Irvine DJ, Purbhoo MA, Krogsgaard M, Davis MM. Direct observation of ligand recognition by T cells. Nature. 2002;419(6909):845–849.

4. Huang J, Brameshuber M, Zeng X, Xie J, Li Qj, Chien Yh, et al. A single peptide-major histocompatibility complex ligand triggers digital cytokine secretion in CD4+ T cells. Immunity. 2013;39(5):846–857.

5. Purbhoo M, Irvine D, Huppa J, Davis M. T cell killing does not require the formation of a stable mature immunological synapse. Nature immunology. 2004;5:524–30. doi:10.1038/ni1058.

6. Stone JD, Chervin AS, Kranz DM. T-cell receptor binding affinities and kinetics: impact on T-cell activity and specificity. Immunology. 2009;126(2):165–176.

7. Govern CC, Paczosa MK, Chakraborty AK, Huseby ES. Fast on-rates allow short dwell time ligands to activate T cells. Proceedings of the National Academy of Sciences. 2010;107(19):8724–8729.

8. Lin JJ, Low-Nam ST, Alfieri KN, McAffee DB, Fay NC, Groves JT. Mapping the stochastic sequence of individual ligand-receptor binding events to cellular activation: T cells act on the rare events. Science Signaling. 2019;12(564):eaat8715.

9. Kersh GJ, Allen PM. Structural Basis for T Cell Recognition of Altered Peptide Ligands: A Single T Cell Receptor Can Productively Recognize a Large Continuum of Related Ligands. J Exp Med. 1996;184(4):1259–68.

10. Kersh GJ, Kersh EN, Fremont DH, Allen PM. High- and Low-Potency Ligands with Similar Affinities for the TCR: The Importance of Kinetics in TCR Signaling. Immunity. 1998;9(6):817–826.

11. Lyons DS, Lieberman SA, Hampl J, Boniface JJ, Chien Yh, Berg LJ, et al. A TCR binds to antagonist ligands with lower affinities and faster dissociation rates than to agonists. Immunity. 1996;5(1):53–61.

12. Altan-Bonnet G, Germain RN. Modeling T cell antigen discrimination based on feedback control of digital ERK responses. PLoS biology. 2005;3(11):e356.

13. Ma Z, Sharp KA, Janmey PA, Finkel TH. Surface-anchored monomeric agonist pMHCs alone trigger TCR with high sensitivity. PLoS biology. 2008;6(2):e43.

14. Feinerman O, Germain RN, Altan-Bonnet G. Quantitative challenges in understanding ligand discrimination by αβ T cells. Molecular immunology. 2008;45(3):619–631.

15. O’Donoghue GP, Pielak RM, Smoligovets AA, Lin JJ, Groves JT. Direct single molecule measurement of TCR triggering by agonist pMHC in living primary T cells. Elife. 2013;2:e00778.

16. FranÇcois P, Voisinne G, Siggia ED, Altan-Bonnet G, Vergassola M. Phenotypic model for early T-cell activation displaying sensitivity, specificity, and antagonism. Proceedings of the National Academy of Sciences. 2013;110(10):E888–E897.

17. Dushek O, van der Merwe PA. An induced rebinding model of antigen discrimination. Trends in immunology. 2014;35(4):153–158.

18. Chakraborty AK, Weiss A. Insights into the initiation of TCR signaling. Nature immunology. 2014;15(9):798–807.

19. Hong J, Ge C, Jothikumar P, Yuan Z, Liu B, Bai K, et al. A TCR mechanotransduction signaling loop induces negative selection in the thymus. Nature immunology. 2018;19(12):1379–1390.

20. Fernandes RA, Ganzinger KA, Tzou JC, Jönsson P, Lee SF, Palayret M, et al. A cell topography-based mechanism for ligand discrimination by the T cell receptor. Proceedings of the National Academy of Sciences. 2019;116(28):14002–14010.

21. Wu P, Zhang T, Liu B, Fei P, Cui L, Qin R, et al. Mechano-regulation of peptide-MHC class I conformations determines TCR antigen recognition. Molecular cell. 2019;73(5):1015–1027.

22. Ganti RS, Lo WL, McAffee DB, Groves JT, Weiss A, Chakraborty AK. How the T cell signaling network processes information to discriminate between self and agonist ligands. Proceedings of the National Academy of Sciences. 2020;117(42):26020–26030.

23. Matsui K, Boniface JJ, Steffner P, Reay PA, Davis MM. Kinetics of T-cell receptor binding to peptide/I-Ek complexes: correlation of the dissociation rate with T-cell responsiveness. Proceedings of the National Academy of Sciences. 1994;91(26):12862–12866.

24. Williams CB, Engle DL, Kersh GJ, Michael White J, Allen PM. A kinetic threshold between negative and positive selection based on the longevity of the T cell receptor–ligand complex. The Journal of experimental medicine. 1999;189(10):1531–1544.

25. Daniels MA, Teixeiro E, Gill J, Hausmann B, Roubaty D, Holmberg K, et al. Thymic selection threshold defined by compartmentalization of Ras/MAPK signalling. Nature. 2006;444(7120):724–729.

26. Corse E, Gottschalk RA, Krogsgaard M, Allison JP. Attenuated T cell responses to a high-potency ligand in vivo. PLoS biology. 2010;8(9):e1000481.

27. Yousefi OS, Günther M, Hörner M, Chalupsky J, Wess M, Brandl SM, et al. Optogenetic control shows that kinetic proofreading regulates the activity of the T cell receptor. Elife. 2019;8:e42475.

28. McKeithan TW. Kinetic proofreading in T-cell receptor signal transduction. Proceedings of the National Academy of Sciences. 1995;92(11):5042.

29. Rabinowitz JD, Beeson C, Lyons DS, Davis MM, McConnell HM. Kinetic discrimination in T-cell activation. Proceedings of the National Academy of Sciences. 1996;93(4):1401–1405. doi:10.1073/pnas.93.4.1401.

30. Chan C, George AJT, Stark J. T cell sensitivity and specificity - kinetic proofreading revisited. Discrete and Continuous Dynamical Systems - B. 2003;3(3):343. doi:10.3934/dcdsb.2003.3.343.

31. Coombs D, Goldstein B. T cell activation: Kinetic proofreading, serial engagement and cell adhesion. Journal of Computational and Applied Mathematics. 2005;184(1):121–139. doi:https://doi.org/10.1016/j.cam.2004.07.035.

32. Jansson A. Kinetic proofreading and the search for nonself-peptides. Self/Nonself. 2011;2(1):1–3. doi:10.4161/self.2.1.15362.

33. Cui W, Mehta P. Identifying feasible operating regimes for early T-cell recognition: The speed, energy, accuracy trade-off in kinetic proofreading and adaptive sorting. PLOS ONE. 2018;13(8):1–16. doi:10.1371/journal.pone.0202331.

34. Ganti RS, Lo WL, McAffee DB, Groves JT, Weiss A, Chakraborty AK. How the T cell signaling network processes information to discriminate between self and agonist ligands. Proc Natl Acad Sci U S A. 2020;117(42):26020–26030.

35. Siller-Farfán JA, Dushek O. Molecular mechanisms of T cell sensitivity to antigen. Immunological reviews. 2018;285(1):194–205.

36. Kirby D, Zilman A. Proofreading does not result in more reliable ligand discrimination in receptor signaling due to its inherent stochasticity. Proceedings of the National Academy of Sciences. 2023;120(21):e2212795120.

37. Huang WYC, Alvarez S, Kondo Y, Lee YK, Chung JK, Lam HYM, et al. A molecular assembly phase transition and kinetic proofreading modulate Ras activation by SOS. Science (American Association for the Advancement of Science). 2019;363(6431):1098–1103.

38. Tischer DK, Weiner OD. Light-based tuning of ligand half-life supports kinetic proofreading model of T cell signaling. Elife. 2019;8:e42498.

39. Pettmann J, Huhn A, Shah EA, Kutuzov MA, Wilson DB, Dustin ML, et al. The discriminatory power of the T cell receptor. eLife. 2021;10:e67092.

40. Bousso P, Robey E. Dynamics of CD8+ T cell priming by dendritic cells in intact lymph nodes. Nature immunology. 2003;4(6):579–585.

41. Miller MJ, Hejazi AS, Wei SH, Cahalan MD, Parker I. T cell repertoire scanning is promoted by dynamic dendritic cell behavior and random T cell motility in the lymph node. Proceedings of the National Academy of Sciences. 2004;101(4):998–1003.

42. Catron DM, Itano AA, Pape KA, Mueller DL, Jenkins MK. Visualizing the first 50 hr of the primary immune response to a soluble antigen. Immunity. 2004;21(3):341–347.

43. Freiberg BA, Kupfer H, Maslanik W, Delli J, Kappler J, Zaller DM, et al. Staging and resetting T cell activation in SMACs. Nature immunology. 2002;3(10):911–917.

44. Celli S, Lemaître F, Bousso P. Real-time manipulation of T cell-dendritic cell interactions in vivo reveals the importance of prolonged contacts for CD4+ T cell activation. Immunity. 2007;27(4):625–634.

45. Bousso P. T-cell activation by dendritic cells in the lymph node: lessons from the movies. Nature Reviews Immunology. 2008;8(9):675–684.

46. Fooksman DR, Vardhana S, Vasiliver-Shamis G, Liese J, Blair DA, Waite J, et al. Functional anatomy of T cell activation and synapse formation. Annual review of immunology. 2009;28:79–105.

47. Wernimont SA, Wiemer AJ, Bennin DA, Monkley SJ, Ludwig T, Critchley DR, et al. Contact-dependent T cell activation and T cell stopping require talin1. The Journal of Immunology. 2011;187(12):6256–6267.

48. Lee KH, Holdorf AD, Dustin ML, Chan AC, Allen PM, Shaw AS. T cell receptor signaling precedes immunological synapse formation. Science. 2002;295(5559):1539–1542.

49. Waldman MM, Rahkola JT, Sigler AL, Chung JW, Willett BAS, Kedl RM, et al. Ena/VASP Protein-Mediated Actin Polymerization Contributes to Naïve CD8+ T Cell Activation and Expansion by Promoting T Cell–APC Interactions In Vivo. Frontiers in Immunology. 2022;13. doi:10.3389/fimmu.2022.856977.

50. Mempel TR, Henrickson SE, Von Andrian UH. T-cell priming by dendritic cells in lymph nodes occurs in three distinct phases. Journal of immunology (Baltimore, Md : 1950). 2004;427(6970):154–159.

51. Henrickson SE, Mempel TR, Mazo IB, Liu B, Artyomov MN, Zheng H, et al. T cell sensing of antigen dose governs interactive behavior with dendritic cells and sets a threshold for T cell activation. Nature immunology. 2008;9(3):282–291.

52. Henrickson SE, Perro M, Loughhead SM, Senman B, Stutte S, Quigley M, et al. Antigen availability determines CD8+ T cell-dendritic cell interaction kinetics and memory fate decisions. Immunity. 2013;39(3):496–507.

53. Le Borgne M, Raju S, Zinselmeyer BH, L. VT, Li J, Wang Y, et al. Real-Time Analysis of Calcium Signals during the Early Phase of T Cell Activation Using a Genetically Encoded Calcium Biosensor. Journal of immunology (Baltimore, Md : 1950). 2016;196(4):1471.

54. Chudnovskiy A, Pasqual G, Victora GD. Studying interactions between dendritic cells and T cells in vivo. Current opinion in immunology. 2019;58:24–30.

55. Desalvo A, Bateman F, James E, Morgan H, Elliott T. Time-resolved microwell cell-pairing array reveals multiple T cell activation profiles. Lab on a Chip. 2020;20(20):3772–3783.

56. Davis DM. Mechanisms and functions for the duration of intercellular contacts made by lymphocytes. Nature Reviews Immunology. 2009;9(8):543–555.

57. Goyette J, Depoil D, Yang Z, Isaacson SA, Allard J, van der Merwe PA, et al. Dephosphorylation accelerates the dissociation of ZAP70 from the T cell receptor. Proceedings of the National Academy of Sciences. 2022;119(9):e2116815119.

58. Shin G, Wang J. The role of energy cost on accuracy, sensitivity, specificity, speed and adaptation of T cell foreign and self recognition. Physical Chemistry Chemical Physics. 2021;23(4):2860–2872.

59. Morgan J, Pettmann J, Dushek O, Lindsay AE. T cell microvilli simulations show operation near packing limit and impact on antigen recognition. Biophys J. 2022;121(21):4128–4136.

60. Coombs D. First among equals: Comment on “Redundancy principle and the role of extreme statistics in molecular and cellular biology” by Z. Schuss, K. Basnayake and D. Holcman. Phys Life Rev. 2019;28:92–93.

61. Schuss Z, Basnayake K, Holcman D. Redundancy principle and the role of extreme statistics in molecular and cellular biology. Physics of Life Reviews. 2019;28:52–79.

62. Lawley SD. Distribution of extreme first passage times of diffusion. Journal of Mathematical Biology. 2020;80(7):2301–2325.

63. Lindsay AE, Bernoff AJ, Navarro Hernández A. Short-time diffusive fluxes over membrane receptors yields the direction of a signalling source. Royal Society Open Science. 2023;10(4):221619.

64. van Oers NS, Killeen N, Welss A. ZAP-70 is constitutively associated with tyrosine-phosphorylated TCR ζ in murine thymocytes and lymph node T cells. Immunity. 1994;1(8):675–685.

65. Kersh GJ, Miley MJ, Nelson CA, Grakoui A, Horvath S, Donermeyer DL, et al. Structural and functional consequences of altering a peptide MHC anchor residue. The Journal of Immunology. 2001;166(5):3345–3354.

66. Love PE, Hayes SM. ITAM-mediated signaling by the T-cell antigen receptor. Cold Spring Harbor perspectives in biology. 2010;2(6):a002485.

67. Lawley SD. Extreme first-passage times for random walks on networks. Phys Rev E. 2020;102:062118. doi:10.1103/PhysRevE.102.062118.

68. Pettmann J, Abu-Shah E, Kutuzov M, Wilson DB, Dustin ML, Davis SJ, et al. T cells exhibit unexpectedly low discriminatory power and can respond to ultra-low affinity peptide-MHC ligands. bioRxiv. 2020;.

69. Voisinne G, Locard-Paulet M, Froment C, Maturin E, Menoita MG, Girard L, et al. Kinetic proofreading through the multi-step activation of the ZAP70 kinase underlies early T cell ligand discrimination. Nature Immunology. 2022;23(9):1355–1364.

