## Supplementary material for "Modulation of antigen discrimination by duration of immune contacts in a kinetic proofreading model of T cell activation with extreme statistics"

### 1 First passage time model of T cell activation

We explore the role of cellular contact time in a model of T cell activation in which a single receptor-ligand complex reaches a signaling state as described by the kinetic proofreading mechanism. In this framework, we derive the distribution of first activation times which gives a fuller interpretation of receptor activation compared to only steady state, or mean, receptor activation concentrations. In addition, the significance of steady state values may change as cellular contact times change, and the extent of this effect is unclear. But in a first passage activation model, changes in cellular contact times would either decrease, or increase, the probability of T cell activation, and the impact of such changes would be clear if the first passage statistics are known.

Understanding the extreme first passage statistics (first passage activation of a large ligand population) may be essential for understanding T cell sensitivity to self antigen. Unlike other processes where extreme statistics offer a biological advantage [1], this is a case in which the T cell must be resistant to the extreme first passage of thousands of self antigen. In the following derivations of the first passage statistics, we assume that self antigen are expressed in much larger quantities than agonist antigen. In addition, we assume that the first passage activation of a receptor by an agonist is not influenced by the self antigen. Conversely, we assume that the first passage activation of a receptor by self antigen is not influenced by the agonist population. Lastly, we assume that if there is an agonist present in the cell contact, then there is only a single agonist present, i.e., the agonist population is always set to 1 or 0. This last assumption is in order to allow the explicit expression of the first passage statistic and to evaluate the more extreme conditions of the cellular contact, since the most difficult task for the model would be to identify a single agonist, when an agonist is present.

#### 1.1 First passage activation of an agonist antigen

We now describe our methodology for determining the full distribution of times for the first complex to reach the signaling state. In the case that there is a single ($n = 1$) ligand and precisely $M$ receptor-ligand dissociation events before an activation event, then the activation time $T_{1,n=1}|M$ has the decomposition

$$T_{1,n=1}|M = t_A + \sum_{m=1}^{M+1} t_B + \sum_{m=1}^{M} t_D. \tag{1}$$

The term $t_A$ is the random variable associated with the observed time that a receptor-ligand complex passes through all bound states without dissociation. Our model assumes that the activated state $C_{n_{KP}}$ is absorbing, therefore the receptor activation event occurs with probability one (given that there are no time constraints), and only occurs once. The distribution for $t_A$ is found by conditioning

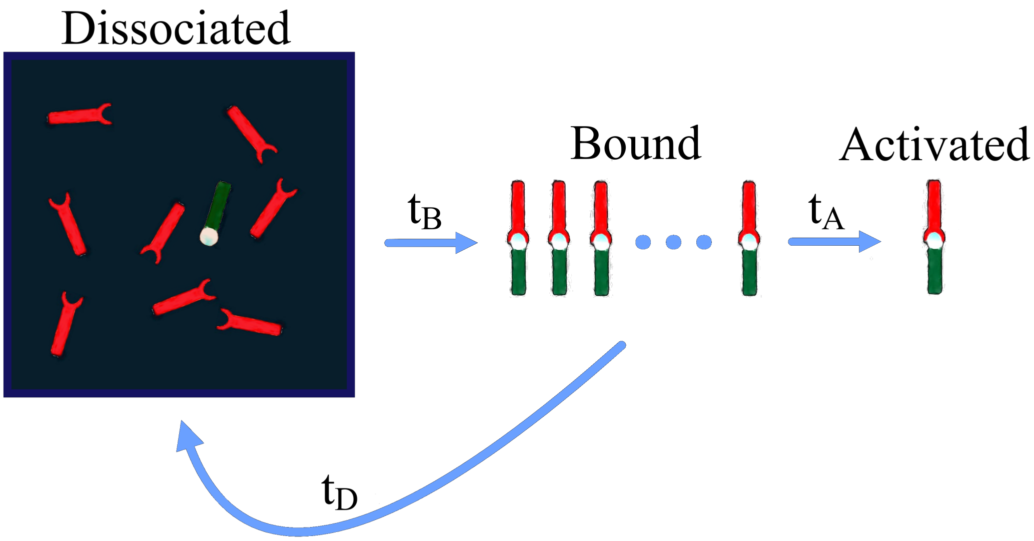


**Fig 1.** The first passage model of a kinetic proofreading mechanism with second order binding and only a single ligand. The first passage time distribution of this model is shown in (13). This model is identical to a model that ignores second order binding. This is because after every dissociation, the population of unbound receptors is the total number of receptors. Thus, every binding event occurs with the same rate, i.e., $K_1 = k_1 R_T$, where $R_T$ is the total number of receptors, for all binding events. The only difference between this model and a model where receptor-ligand pairs are independent is an increased on-rate due to multiple receptor influence on binding events.

on another random variable $t_{n_{KP}}$, which is the time it takes for a bound receptor-ligand pair to become activated given that dissociation is not possible, and the random variable $t_{-1}$, which is the time it takes for a receptor-ligand pair to dissociate given that there is no absorbing state. The unconditioned random variables are

$$t_{n_{KP}} \sim \text{Erlang}(n_{KP}, k_p), \qquad t_{-1} \sim \text{Exp}(k_{-1}), \tag{2}$$

and the conditioned variable is

$$t_A = \{t_{n_{KP}} | t_{n_{KP}} < t_{-1}\}. \tag{3}$$

In Eq. (2) the Erlang$(k, \lambda)$ distribution describes the distribution of a sum of $k$ independent exponential variables of common mean $\lambda^{-1}$ and has distribution

$$f_E(t; k, \lambda) = \frac{\lambda^k t^{k-1} e^{-\lambda t}}{(k-1)!}.$$

The density of $t_A$, $f_A(t)$, is found by computing

$$\begin{aligned} f_A(t) = \mathbb{P}(t_{n_{KP}} = t | t < t_{-1}) &= \frac{\mathbb{P}(t_{-1} > t | t_{n_{KP}} = t)\mathbb{P}(t_{n_{KP}} = t)}{\int_0^\infty \mathbb{P}(t_{-1} > t | t_{n_{KP}} = t)\mathbb{P}(t_{n_{KP}} = t) dt} \\ &= \frac{e^{-(k_{-1}+k_p)t}(k_{-1} + k_p)^{n_{KP}} t^{n_{KP}-1}}{(n_{KP} - 1)!}. \end{aligned} \tag{4}$$

Here $t_A \sim$ Erlang$(n_{KP}, k_p + k_{-1})$. The random variable, $t_B$, is the time a receptor-ligand pair spends in an unbound state, hence,

$$t_1 \sim \text{Exp}(K_1), \qquad f_B(t) = K_1 e^{-K_1 t}. \tag{5}$$

The random variable, $t_D$, is the observed dissociation time given that this dissociation event occurred before the receptor-ligand complex reached the activated state. Similarly to $t_A$, this is found by conditioning on $t_{-1}$ and $t_{n_{KP}}$. The conditioned random variable is then

$$t_D = \{t_{-1} | t_{-1} < t_{n_{KP}}\},$$

where the PDF is found by the method above to be

$$f_D(t) = \mathbb{P}(t_{-1} = t | t < t_{n_{KP}}) = \frac{\mathbb{P}(t_{n_{KP}} > t | t_{-1} = t)\mathbb{P}(t_{-1} = t)}{\int_0^\infty \mathbb{P}(t_{n_{KP}} > t | t_{-1} = t)\mathbb{P}(t_{-1} = t)dt} = \frac{e^{-k_{-1}t} k_{-1} \Gamma(n_{KP}, k_p t)}{(n_{KP} - 1)!(1 - \alpha^n_{KP})}.$$

Here $\Gamma(n_{KP}, k_p t)$ is the upper incomplete gamma function. Lastly, we have the random variable, M, which is the number of dissociation events that occur before a receptor-ligand pair reaches the activated state. This is equivalent to counting the number of failures before a success occurs. Thus,

$$M \sim \text{Geometric}(\alpha^{n_{KP}}), \qquad \alpha = \frac{k_p}{k_p + k_{-1}}, \qquad \mathbb{P}(M = m) = \alpha^{n_{KP}}(1 - \alpha^{n_{KP}})^m, \tag{6}$$

where $n_{KP}$ is the number of kinetic proofreading steps. This formulation leads to (1), which is a sum of independent random variables. To find $f_T(t)$, we utilize the properties of the associated characteristic functions,

$$\psi_A(x) = \int_{-\infty}^{\infty} e^{-ixt} f_A(t)dt, \quad \psi_B(x) = \int_{-\infty}^{\infty} e^{-ixt} f_B(t)dt, \quad \psi_D(x) = \int_{-\infty}^{\infty} e^{-ixt} f_D(t)dt. \tag{7}$$

The summations in (1) then becomes

$$\mathbb{E}[e^{ix\sum_{m=1}^{M+1} t_B}] = \psi_B^{M+1}, \qquad \mathbb{E}[e^{ix\sum_{m=1}^{M} t_D}] = \psi_D^M. \tag{8}$$

Putting the terms together gives the characteristic function of the conditional distribution of T.

$$\psi_{T|M=m} = \psi_A \psi_B^{m+1} \psi_D^m. \tag{9}$$

The random variable, $T|M = m$, is the distribution of the first passage time given that there were m dissociation events (or equivalently: m failures). We know that the sequence of events, $\{T|M = m\}_{m=1}^{\infty}$, form a disjoint partition of the sample space. From an observers point of view, the probability we "observe" a random variable with characteristic function $\psi_{T|M=m}$ is $(1-\alpha^{n_{KP}})^m \alpha^{n_{KP}}$. Thus, the characteristic function of the full distribution $\Psi_T$ can be decomposed into a sum of conditioned characteristic functions, i.e.,

$$\psi_T = \sum_{m=1}^{\infty} \psi_A \psi_B^{m+1} \psi_D^m (1 - \alpha^{n_{KP}})^m \alpha^{n_{KP}}, \tag{10}$$

replacing the respective characteristic functions with their associated parameters yields the explicit characteristic function of the first passage time model for a single ligand,

$$\psi_T = \frac{K_1 k_p^{n_{KP}} (ik_{-1} + x)}{(K_1 - ix)(ik_{-1} k_p^{n_{KP}} + (k_{-1} + k_p - ix)^{n_{KP}} x)}. \tag{11}$$

Finally, to obtain the distribution of the first passage time for a kinetic proofreading model with only first order reactions, we obtain the inverse characteristic integral

$$f_T(t) = \frac{1}{2\pi} \int_{-\infty}^{\infty} e^{-ixt} \frac{K_1 k_p^{n_{KP}} (ik_{-1} + x)}{(K_1 - ix)(ik_{-1} k_p^{n_{KP}} + (k_{-1} + k_p - ix)^{n_{KP}} x)} dx \tag{12}$$

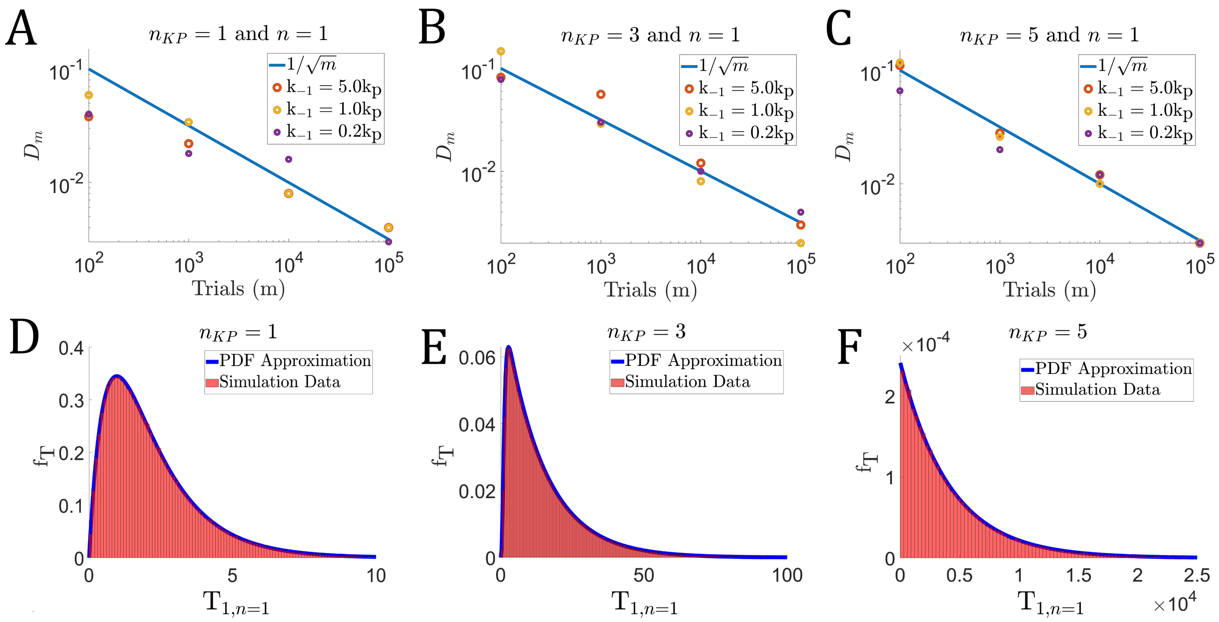


**Fig 2. A–C** The Kolmogorov-Smirnov statistic for models with $n_{KP}$=1, $n_{KP}$=3, and $n_{KP}$=5 kinetic proofreading steps. Three dissociation rates were used and described in reference to their relationship to the forward rate, $k_p$, with $k_p = 1s^{-1}$. A K-S statistic was computed with different numbers of simulated first passage activation trials (m) for each dissociation rate. The number of trials were increased 10-fold for each measure, and there is a clear observable decrease in the statistic for each of the models shown, indicating a possible convergence to the CDF of (13). **D–F** Plots showing (13) with histograms obtained from stochastic simulation using the Gillespie Algorithm. The first plot, **D**, shows a data from a model with a single kinetic proofreading step and $k_{-1} = 0.2k_p$. Plot **E** shows a model with $n_{KP}$=3 and $k_{-1} = 1.0k_p$. Lastly, **F** shows a model with $n_{KP}$=5 kinetic proofreading steps and $k_{-1} = 5.0k_p$. From left to right, **D–F** shows data from models in which the first passage activation time is increasing and $\alpha^{n_{KP}}$ is decreasing.

The integral (13) is readily evaluated numerically. To demonstrate the accuracy of (13), we compare the analytical distribution (13) with stochastic simulations. For comparison, we utilize the Kolmogorov-Smirnov (K-S) test statistic

$$D_m = \sup_{t>0} |F_m(t) - F(t)|. \tag{13}$$

where $F_m(t)$ is the empirical CDF obtained from stochastic simulation of the first passage activation model. In Figure 2A–C, we demonstrate the accuracy of (13) by observing the K-S statistic decrease to zero as the number of stochastic trials $(m)$ increases. In addition, we show in Figure 2 D–F that the analytical PDF (13) agrees very well with the first passage activation data obtained from stochastic simulation. We observe that the character of the distribution is dynamic with respect to model parameters, with an exponential behavior for larger $k_{-1}$ values and more kinetic proofreading steps $(n_{KP})$, while it assumes a gamma distribution when $k_{-1}$ is small and there are only a few kinetic proofreading steps. This structural sensitivity to parameters variations has previously been observed in [2].

#### 1.2 First passage receptor activation of a large antigen population

In the scenario of a large number of receptors and ligands, finding an analytical expression for the first passage time is more challenging to obtain due to the the nonlinear binding rate $K_1(R(t), L(t))$. Here we propose a simple extension to the KP system of equations to obtain an approximation of the extreme statistics. We first define $A(t)$ to be the probability that a T cell is non-activated by time $t$ (CDF). We introduce a modification of the ODE system that describes the conditioned dynamics of non-activated receptors, i.e. those that have not have entered the activated state ($C_{n_{KP}}$) by time $t$. Their dynamics is governed by the ODE system

$$\frac{dR}{dt} = k_{-1} \sum_{i=0}^{n_{KP}-1} C_i + k_p C_{n_{KP}-1} - k_1 R(L_T - (R_T - R)) \tag{14a}$$

$$\frac{dC_0}{dt} = k_1 R(L_T - (R_T - R)) - (k_{-1} + k_p) C_0 \tag{14b}$$

$$\vdots$$

$$\frac{dC_i}{dt} = k_p C_{i-1} - (k_{-1} + k_p) C_i \tag{14c}$$

$$\vdots$$

$$\frac{dC_{n_{KP}-1}}{dt} = k_p C_{n_{KP}-2} - (k_{-1} + k_p) C_{n_{KP}-1}. \tag{14d}$$

This system features an adjustment to equation (14a). The dynamics of $A(t)$ then follows

$$\frac{dA}{dt} = k_p C_{n_{KP}-1}(1 - A). \tag{14e}$$

For example, in the model described in Figure 1, the bound population in $C_{n_{KP}-1}(t)$ can be interpreted as the population in $C_{n_{KP}-1}$ at time $t$ given that no receptors have been activated. However, since the dwell time in state $C_{n_{KP}-1}$ is distributed according to $\text{Exp}(k_{-1} + k_p)$, and the activated state ($C_{n_{KP}}$) is an absorbing state, then we assume that $k_{-1} + k_p$ is the rate of "leaving" state $C_{n_{KP}-1}$ in our conditioned system.

Here $R(t)$ is the unbound receptor population at time $t$, $R_T$ and $L_T$ are the conserved total population of receptors and ligands, respectively, and $L(t) = L_T - (R_T - R)$ is the population of unbound ligands. In (14e), $A = A(t)$, with $A(0) = 0$, is the probability that the kinetic proofreading system has produced an activated receptor by time $t$ such that when (14) is solved numerically, the function $A = A(t)$ becomes the approximation to the CDF of the extreme statistic.

This assumption is one source of error in this approximation. The other source of error is the assumption the population is continuous, since we are utilizing a continuous population model to describe a discrete stochastic process. However, the magnitude of both errors decrease as $\alpha_{KP}^n \to 0$ and the population increases (Figure 4A–F). Furthermore, the error in our approximation of the conditioned dissociation rate vanishes when $n_{KP} = 1$. In Figure 4, we show the K-S statistic for several cases as well as compare the probability density obtained from the ODE system ($f_{\text{T}} = dA/dt$) with that of data obtained from stochastic simulation.

When there is only a single kinetic proofreading step, $n_{KP} = 1$, the above ODE system becoming the true conditioned mass action model (as opposed to an approximation). The error in our ODE approximation method is that we assume that the only change in dissociation rate occurs at the state $C_{n_{KP}-1}$ (Figure 3). In reality each of the dissociation rates of the system is affected when the mass action kinetics are "conditioned" so. However, when $n_{KP} = 1$, the distribution of the dissociation

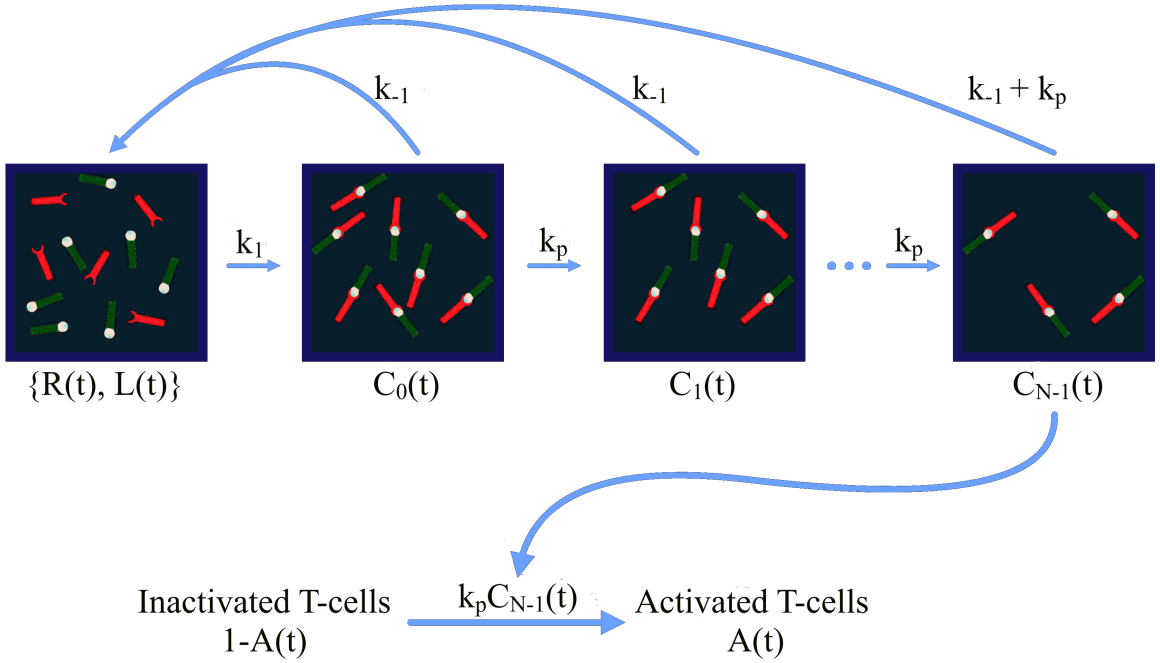


**Fig 3.** A diagram of the $n_{KP}$+2 state ODE model. Each state, with the exception of $C_{n_{KP}}$, evolves continuously over time. The difference between this model and the traditional mass action model is that this system is evolved with the condition that no TCRs have been activated. The state, A(t), is the fraction of activated systems by time $t$, i.e., the fractions of systems that have produced an activated TCR. This is also an approximation of the CDF for the first passage time for system activation.

time, $t_D$, becomes an exponential distribution, i.e., $t_D \sim \text{Exp}(k_p + k_{-1})$. Unlike the distribution, $f_D$, found above, this yields a random variable for which the asymptotics are easily computed. Now, dissociation in the ODE approximation exactly matches that of the asymptotics $(k_p + k_{-1})$ of the conditioned dissociation time, $t_D$. In this case, the approximate conditioned mass action system becomes the exact conditioned mass action system, for the first passage activation model. When $n_{KP} > 1$, the approximation of the dissociation rates do not exactly match the asymptotics of the conditioned dissociation time, $t_D$. But, these differences decrease as the probability of activation decreases. This is because the conditioned dissociation time, $t_D$, is conditioned such that dissociation occurred faster than activation of a receptor ligand complex, $t_{-1} < t_{n_{KP}}$. As the dissociation rate grows larger than the forward rate, $k_{-1} \gg k_p$, or the number of kinetic proofreading steps increase, then $(1 - \alpha^{n_{KP}}) = \mathbb{P}(t_{-1} < t_{n_{KP}}) \to 1$. In this case, the conditioned mass action system behaves nearly identically to the unconditioned mass action system, i.e., the mass action system where dissociation from each complex state, $C_i$, occurs with rate $k_{-1}$. One can see this by setting $k_p = k_{-1}/\kappa$ and taking the limit as $\kappa \to \infty$ in the above description for $f_D$, which yields $f_D \to k_{-1}e^{-k_{-1}t}$. Changing our ODE approximation to reflect this yields an accurate approximation to the first passage activation time in this case.

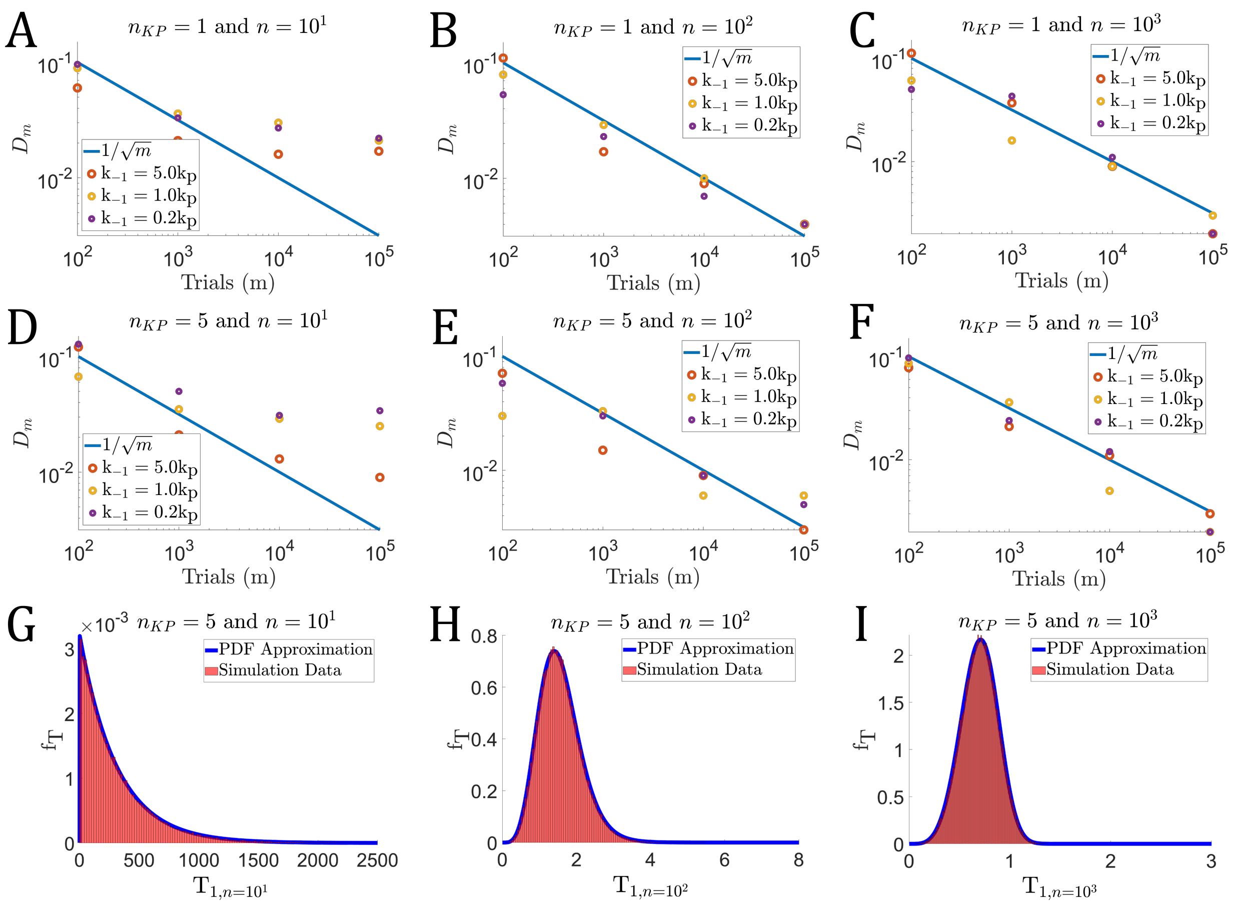

**Fig 4.** **A–F** Convergence of approximate ODE system (14) to stochastic simulations The Kolmogorov-Smirnov statistic for models with $n_{KP}$=1 and $n_{KP}$=5 kinetic proofreading steps and ligand population n. In the cases shown, the total number of receptors is equal to the number of ligands. $R_T = L_T = n$. Three dissociation rates were used and described in reference to their relationship to the forward rate, $k_p$, with $k_p = 1s^{-1}$. Similar to Figure 2, K-S statistic was computed with different numbers of simulated first passage activation trials (m) for each dissociation rate. However, unlike in Figure 2, some cases do not yield a decrease in the K-S statistic even given a 10-fold increase in the number of trials (**A** and **D**). The primary reason for this is that the approximation assumes continuous populations, but the model is of a discrete stochastic process. When the ligand population is increased, we observe similar behavior in the K-S statistic as in the single ligand case (**B**, **C**, **E** and **F**). **G–I** Plots showing the ODE solution ($f_T = dA/dt$) with histograms obtained from stochastic simulation using the Gillespie Algorithm. Each plot shows results from a model with 5 kinetic proofreading steps ($n_{KP} = 5$). **G**: rates $k_{-1} = 5.0k_p$ and $n = 10$. **H**: rates $k_{-1} = 1.0k_p$ and $n = 100$. **I**: data with parameters $k_{-1} = 0.2k_p$ and $n = 1000$. **G–I**: first passage activation distributions for $n = 10, 100, 1000$.

#### 2 Limiting behavior of extreme statistics and ODE approximation

Recent work has shown that a continuous time Markov Chain with discrete states converges to a Weibull distribution limit $n \to \infty$ [3]. More specifically, given an extreme arrival time such that

$$\mathrm{T}_{1,n} := \min\{t_1, t_2, \ldots, t_n\}$$

where $\{t_i\}_{i=1}^n$ are $n$ i.i.d. first passage times, then

$$\mathrm{T}_{1,n} \approx_d \text{Weibull}((An)^{-1/k}, k).$$

In other words, $(An)^{-1/k}\mathrm{T}_{1,n}$ converges in distribution to Weibull$(1, k)$. A random variable $X \geq 0$ has a *Weibull* distribution with scale parameter $\lambda > 0$ and shape parameter $k > 0$ if $\mathbb{P}(X > x) = e^{-(x/\lambda)^k}$. In such a case, we define $X =_d$ Weibull$(\lambda, k)$.

In the context of the FRAM, $A = k_p^{n_{KP}}/n_{KP}!$, $\lambda = (An)^{-1/k}$ and $k = n_{KP}$. A source of numerical error in applying this result [3] to a large antigen population in the FRAM is that the first passage activation of receptors are not i.i.d., since the propensity for binding events is $K_{on} = k_{on}RL$. However, we still observe convergence with sufficiently large $n$ (Figure 5). This is due to large receptor and ligand populations effectively reducing the binding times of the fastest first receptor activation time to unobservable values. Finally, it should be noted that since we observed the empirical CDF from stochastic data converges to the ODE approximation (Figure 4) when $n$ is large, then it follows that the empirical distribution obtained from stochastic simulation also converges to a Weibull$((An)^{-1/k}, k)$ when $n$ is sufficiently large.

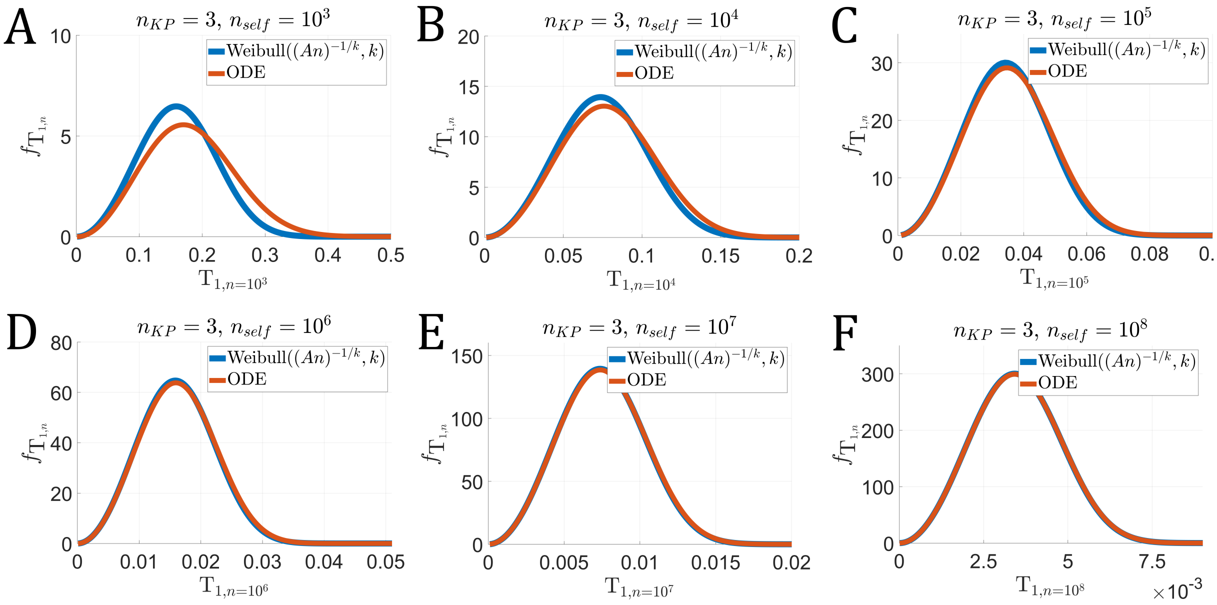


**Fig 5.** Plots showing that the extreme arrival time of a large antigen population ($n = n_{self}$) in the FRAM converges to the result in [3], given a sufficiently large number of receptor-ligand pairs.

#### 3 Random contact times

We modify our first passage activation model to incorporate random cell contact times, as this is a more feasible reality for T cells [4]. Furthermore, utilizing a distribution of contact durations versus that of a single fixed time demonstrates the robustness of the FRAM with respect to variability in the T cell/APC contact duration. The extension requires little modification to our previous probabilistic descriptions. The primary difference is that $\tau$ is now a random variable, which we arbitrarily choose to be exponentially distributed, i.e., $\tau \sim \text{Exp}(\lambda)$. However, a similar analysis could be performed with any temporal distribution. Thus, for any contact duration $\tau$, we must multiply the outcome probabilities of a contact duration $\tau$ by the probability that the cell contact ends at time $\tau$. Given $f(\tau;\lambda)$, the PDF of $\tau$ given exponential distribution parameter $\lambda$, the outcome probabilities become

$$\mathbb{P}(\text{TP};\lambda) = \int_0^\infty f(\tau;\lambda)\mathbb{P}\left(\text{TP};\tau\right) d\tau$$
$$\mathbb{P}(\text{FP};\lambda) = \int_0^\infty f(\tau;\lambda)\mathbb{P}\left(\text{FP};\tau\right) d\tau$$

Similar to the analysis of a fixed contact duration, we show the outcome probabilities for several different models. However, instead of showing the outcome probabilities with respect to a varying $\tau$, we now show the outcome probabilities over a varying distribution parameter $\lambda$ (Figure 6).

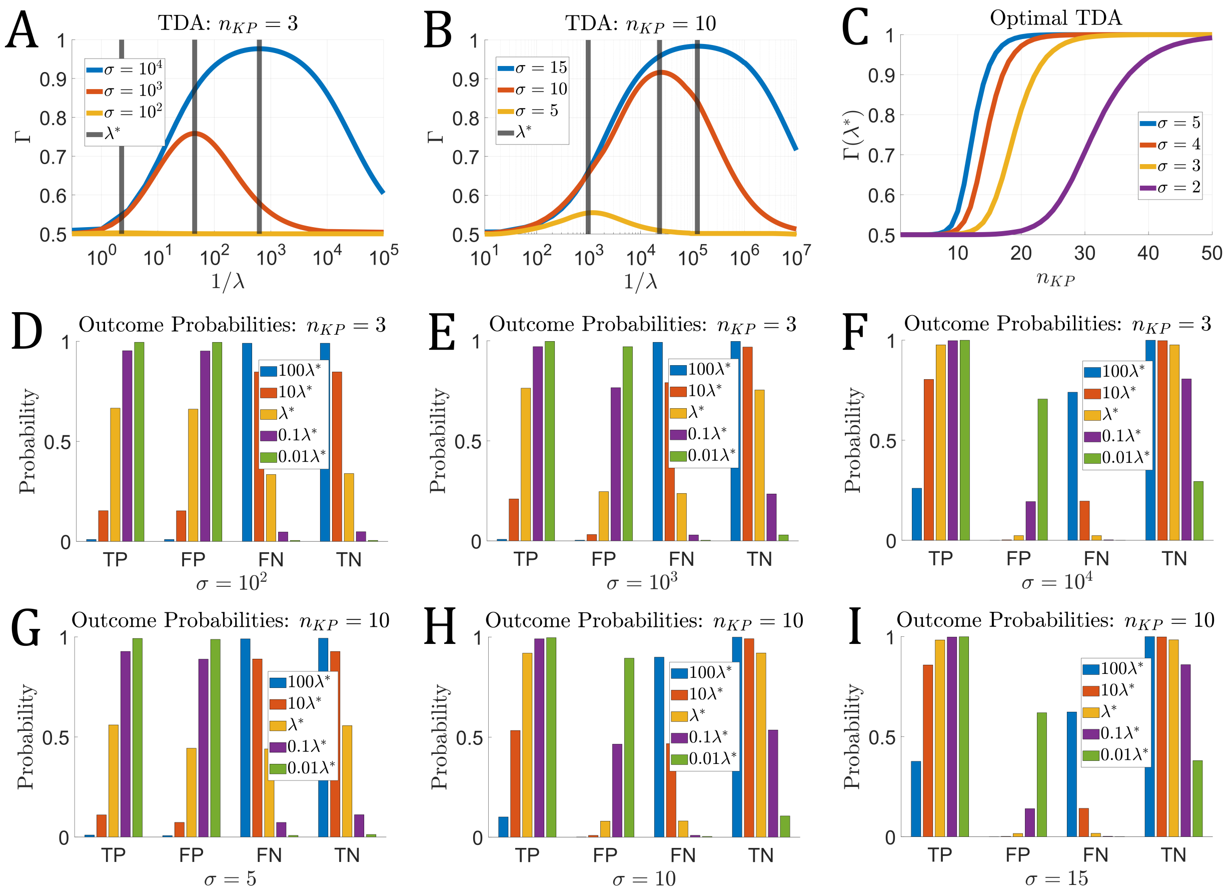


**Fig 6.** The T cell Accuracy $\Gamma(\lambda)$, where the contact time $\tau \sim \text{Exp}(\lambda)$. **A–B** The T cell Accuracy for an $n_{KP} = 3$ and $n_{KP} = 10$ kinetic proofreading model with contact time distribution parameter $\lambda$. Here, as the TDA improves in a model ($\Gamma(\lambda^*)$ increases), the mean contact time also increases ($\lambda$ decreases). **C** As discussed in the fixed contact time case, this also results in a larger number of reactions and longer cellular contact durations. **D–F** Outcome probabilities for a $n_{KP}$=3-step kinetic proofreading model where cellular contact times are exponentially distributed with parameter $\lambda$. **G–I** Outcome Probabilities for a $n_{KP} = 10$-step kinetic proofreading model with exponentially distributed cellular contact durations.

Similar to the case of a fixed contact duration, we note that there is an optimal distribution parameter, $\lambda^*$, for which $\Gamma$ is maximized (Figure 6A–B). Another similarity to that of the fixed contact durations, it is possible to obtain near perfect ligand discrimination given only small differences by increasing the number of kinetic proofreading steps (Figure 6C). It is also true that the optimal parameter, $\lambda^*$, may not be the best strategy for a T cell (Figure 6D–I), due to the same reasons previously mentioned in the fixed contact time case. One difference worth noting is that there is a slight loss of ligand discrimination when $\tau$ is exponentially distributed. If $\Gamma(\lambda^*) = \Gamma_{\lambda^*}$ and $\Gamma(\tau^*) = \Gamma_{\tau^*}$, then the T cell accuracy for the exponentially distributed contact durations is such that $\Gamma_{\lambda^*} \leq \Gamma_{\tau^*}$, for any two identical models. This is because we found the optimal contact duration when time was fixed, which was $\tau^*$. Therefore, any shorter or longer contact durations are suboptimal and will yield a smaller TDA than $(\tau^*)$. Therefore, the best possible distribution for cellular contact durations in the cases where the optimum is unique are those in which $\mathbb{P}(\tau = \tau^*) = 1$, for a random variable $\tau$.

### 4 Time as a cost in the FRAM

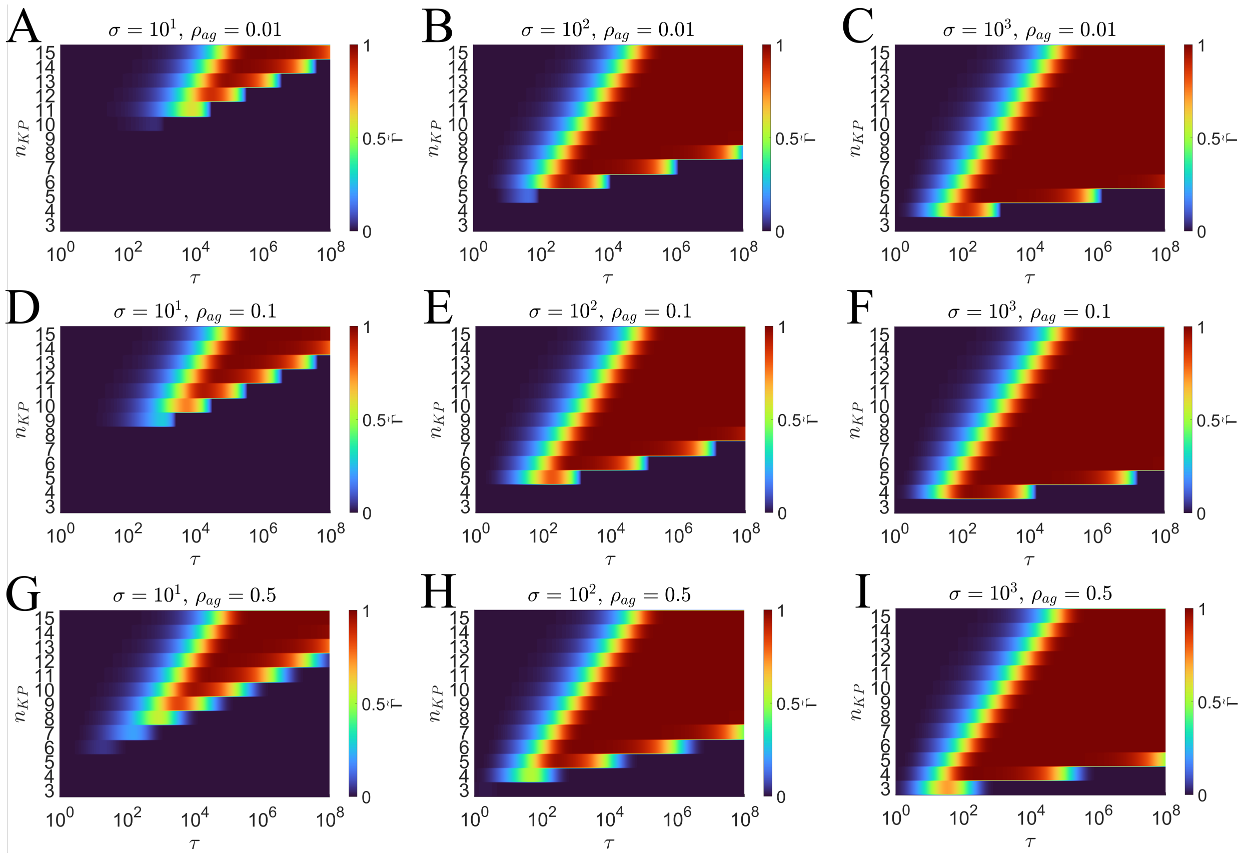


**Fig 7.** Heatmaps showing the cost of high accuracy in the FRAM with respect to the cellular contact duration $\tau$ and $n_{KP}$. The scaled accuracy $\tilde{\Gamma}$ is such that $\tilde{\Gamma} = 0$ is $\Gamma \leq \Gamma^{(BL)}$ and $\tilde{\Gamma} = (\Gamma - \Gamma^{(BL)})/(1 - \Gamma^{(BL)})$. The left, center, and right columns of plots show results for $\sigma = 10$, $\sigma = 10^2$, $\sigma = 10^3$, respectively. Each top, center, and bottom row shows results for $\rho_{ag} = 0.01$, $\rho_{ag} = 0.1$, and $\rho_{ag} = 0.5$.

Assuming the KP rates are fixed, then increases in the number of KP steps will delay the first passage of a receptor triggered by an agonist antigen. Because of this, longer cellular contact durations are necessary in order for a high true positive probability. In Figure 7, we show that challenging classification environments yield longer optimal contact durations. More specifically, as $\sigma$ is decreased (columns in Figure 7), self and agonist antigen become more similar. This challenge is overcome by introducing more kinetic proofreading steps which are needed to effectively distinguish the self and agonist antigen populations in cellular contacts. However, this also results in longer optimal cellular contact durations since the more KP steps means longer wait times for receptor triggering by agonists. Similarly, decreasing the agonist positive prevalence ($\rho_{ag}$) increases the challenge of the environment since agonist positive cells become more rare. The rarity of agonist positive cells means that there may be many agonist negative contacts formed before an agonist positive cell is found. Because of this, even low probabilities of false positives become significant. As before, this challenge is overcome by introducing larger $n_{KP}$, which results in more time.

#### 5 Scaling contact durations in the FRAM

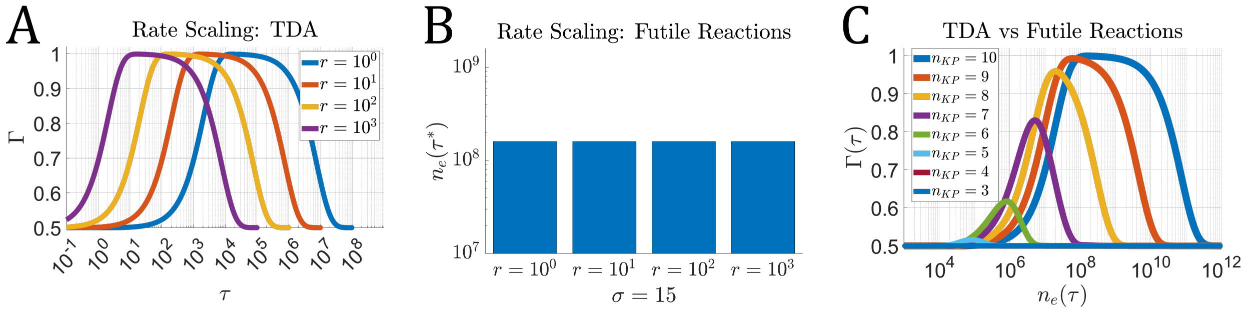


**Fig 8. A** The T cell accuracy $\Gamma$ versus cellular contact duration $\tau$ for $\sigma = 15$ and $n_{KP} = 10$. The scaling $r$ is applied uniformly to all KP rates. In log-space, this results in a translation of the accuracy with respect to time. **B** The mean number of futile reactions at optimal contact durations $n_e(\tau^*)$ for multiple scalings $r$. **C** The accuracy $\Gamma(\tau)$ versus the mean number of futile reactions $n_e(\tau)$ for varying $\tau$ and multiple $n_{KP}$.

In the FRAM, we can scale the optimal contact duration to any value of our choosing without reducing the maximum accuracy. To demonstrate this, we introduce the scaling factor, $r \in \mathbb{R}^+$, such that for the vector of parameters $\theta = (k_1, k_{-1}, k_p)$

$$\theta_r = r\theta = (rk_1, rk_{-1}, rk_p).$$

If $\tau^*$ is the optimal contact duration that maximizes the accuracy in a model with rates $\theta$, then $\tau^*/r$ is the optimal contact duration that maximizes the accuracy in a model with rates $\theta_r$. In fact, the accuracy is identical at every scaled $\tau$, i.e., $\Gamma(\tau/r; \theta_r) = \Gamma(\tau; r)$ (Figure 8A). Thus, any FRAM can be scaled to yield an identical classification accuracy in shorter-duration contacts by scaling the KP reaction rates. However, the number of futile reactions will remain constant with respect to this scaling (Figure 8B). This is due to the temporal evolution of the KP mass action kinetics. By equally increasing all KP rates, the mass action system evolves faster in time. However, the evolution is identical to the model with the original rates, with respect to the scaling, i.e., the state of the KP model with scaled rates at $\tau/r$ is identical to that of the state of the model with the original KP rates at time $\tau$.

#### References


1. Schuss Z, Basnayake K, Holcman D. Redundancy principle and the role of extreme statistics in molecular and cellular biology. Physics of Life Reviews. 2019;28:52–79.

2. Bel G, Munsky B, Nemenman I. The simplicity of completion time distributions for common complex biochemical processes. Physical biology. 2009;7(1):016003.

3. Lawley SD. Extreme first-passage times for random walks on networks. Phys Rev E. 2020;102:062118. doi:10.1103/PhysRevE.102.062118.

4. Cai E, Marchuk K, Beemiller P, Beppler C, Rubashkin MG, Weaver VM, et al. Visualizing dynamic microvillar search and stabilization during ligand detection by T cells. Science. 2017;356(6338):eaal3118.
